# Evolution of the insecticide target *Rdl* in African *Anopheles* is driven by interspecific and interkaryotypic introgression

**DOI:** 10.1101/2019.12.17.879775

**Authors:** Xavier Grau-Bové, Sean Tomlinson, Andrias O. O’Reilly, Nicholas J. Harding, Alistair Miles, Dominic Kwiatkowski, Martin J. Donnelly, David Weetman, The Anopheles gambiae 1000 Genomes Consortium

## Abstract

The evolution of insecticide resistance mechanisms in natural populations of *Anopheles* malaria vectors is a major public health concern across Africa. Using genome sequence data, we study the evolution of resistance mutations in the *resistance to dieldrin locus* (*Rdl*), a GABA receptor targeted by several insecticides, but most notably by the long-discontinued cyclodiene, dieldrin. The two *Rdl* resistance mutations (*296G* and *296S*) spread across West and Central African *Anopheles* via two independent hard selective sweeps that included likely compensatory nearby mutations, and were followed by a rare combination of introgression across species (from *A. gambiae* and *A. arabiensis* to *A. coluzzii*) and across non-concordant karyotypes of the 2La chromosomal inversion. *Rdl* resistance evolved in the 1950s as the first known adaptation to a large-scale insecticide-based intervention, but the evolutionary lessons from this system highlight contemporary and future dangers for management strategies designed to combat development of resistance in malaria vectors.

## Introduction

The recurrent evolution of insecticide resistance in the highly-variable genomes of *Anopheles* mosquitoes (Neafsey et al. 2015; Miles et al. 2017; Clarkson et al. 2019) is a major impediment to the ongoing efforts to control malaria vector populations. Resistance to dieldrin was the first iteration of this cyclical challenge: this organochlorine insecticide was employed in a pioneering vector control programme in Nigeria in 1954, but resistant *Anopheles* had already appeared after just 18 months (Elliott and Ramakrishna 1956) due to a single dominant mutation (Davidson 1956; Davidson and Hamon 1962). Dieldrin use ceased in the 1970s due to its high persistence as an organic pollutant and unexpectedly wide toxicity, culminating in a ban by the 2001 Stockholm Convention on Persistent Organic Pollutants. However, resistance has remained strikingly persistent in natural *Anopheles* populations for more than 40 years (Du et al. 2005). The study of the genetic architecture of dieldrin resistance can thus provide key insights into the evolutionary ‘afterlife’ of resistance mechanisms to legacy insecticides. We address this issue by studying its emergence and dissemination in contemporary African populations of the *A. gambiae* species complex.

Dieldrin resistance in *Anopheles spp.* is caused by mutations in its target site, the γ-aminobutyric-aminobutyric acid (GABA) receptor gene, a ligand-gated chloride channel also known as *resistance to dieldrin locus*—or *Rdl—*that is strongly conserved in a wide range of insects (ffrench-Constant, Rocheleau, et al. 1993; Thompson et al. 1993; Du et al. 2005). Two resistance mutations have been found in anophelines, both in codon 296: alanine-to-glycine (*A296G*) and alanine-to-serine (*A296S*). Resistant mutations in the homologous *Rdl* codon have also evolved in other insects, e.g. in *Drosophila* spp. (codon 302) (ffrench-Constant, Rocheleau, et al. 1993; Thompson et al. 1993; Du et al. 2005). Populations of *Anopheles gambiae* sensu stricto (henceforth, *A. gambiae*) and its sister species *A. coluzzii* possess both *296G* and *296S* alleles (Du et al. 2005; Lawniczak et al. 2010), whereas the *296S* allele is the only one reported in *A. arabiensis* and the more distantly-related malaria vectors *A. funestus* and *A. sinensis* (Du et al. 2005; Wondji et al. 2011; Yang et al. 2017). Normally, dieldrin inhibits the activity of *Rdl* receptors, causing persistent neuronal excitation and rapid death; but codon 296 mutations confer resistance by reducing its sensitivity to the insecticide (ffrench-Constant et al. 2000). However, in the absence of exposure, *Rdl* mutations appear to carry fitness costs, such as lower mosquito mating success (Platt et al. 2015) or impaired response to oviposition and predation-risk signals (Rowland 1991a; Rowland 1991b) (although see (ffrench-Constant and Bass 2017)). Consequently, with seemingly limited current benefit via exposure to insecticides targeting *Rdl*, persistence of the mutations in anophelines is puzzling.

We interrogate the *Anopheles gambiae* 1000 Genomes cohort (The *Anopheles gambiae* 1000 Genomes Consortium 2017; Clarkson et al. 2019) to ascertain how often dieldrin resistance mutations have evolved in the *A. gambiae*/*A. coluzzii* species pair, and the mechanisms by which these alleles spread across Africa and may persist. We identify two distinct *Rdl* resistance haplotypes in these species, defined by hard selective sweeps and the perfect linkage of the *296G* and *296S* alleles with putatively compensatory mutations. Furthermore, the resistance haplotypes are across genomes from different species (*A. gambiae*, *A. coluzzii* and *A. arabiensis*), and across chromosomes with differing karyotypes in the 2La inversion (the longest inversion in *Anopheles* genomes) (Coluzzi 2002) within which *Rdl* resides. Inter-species reproductive isolation and inversions such as 2La both result in reduced recombination rates (Sturtevant 1917; Andolfatto et al. 2001; Ayala and Coluzzi 2005; Kirkpatrick 2010), which would in principle hinder the spread of these adaptive alleles. Here, we provide evidence that *Rdl* resistance alleles, which our structural modelling shows have divergent effects on the channel pore, underwent a rare combination of interspecific and interkaryotypic introgression.

Overall, we show that two founding resistance mutations spread with remarkable ease across geographical distance, species, and recombination barriers. This evolutionary trajectory has parallels with later-emerging target site resistance mutations, such as *Vgsc* (Martinez-Torres et al. 1998; Davies et al. 2007; Clarkson et al. 2014; Clarkson et al. 2018). The persistence of dieldrin resistance also challenges the efficacy of current and newly developed insecticides that also target *Rdl* (Gant et al. 1998; Nakao and Banba 2015; Miglianico et al. 2018), as well as the efficacy of rotative insecticide management strategies (World Health Organization 2012). These results thus emphasise the influence of past interventions on current and future programmes of vector population control.

## Results

### Distribution of *Rdl* resistance mutations across African populations

First, we investigated the genetic variation in *Rdl* across populations of the *Anopheles gambiae* species complex, including *A. gambiae* and *A. coluzzii* from the *Anopheles gambiae* 1000 genomes project (*Ag1000G* Phase 2, *n* = 1142) (Clarkson et al. 2019), and outgroups from four other species (*A. arabiensis*, *A. quadriannulatus*, *A. melas* and *A. merus*; *n* = 36) (Fontaine et al. 2015). All genomes and their populations of origin are listed in Supplementary Material SM1.

We identified six non-synonymous mutations that are segregating in at least one population at ≥5% frequency (Figure 1A; complete list of variants in Supplementary Material SM2), including the *296G* and *296S* resistance alleles. *296G* is present in West and Central African populations of both *A. gambiae* and *A. coluzzii*, with frequencies ranging from 30% (Cameroon *A. gambiae*) to 96% (Ghana *A. gambiae*). *296S* is present in *A. coluzzii* specimens from Burkina Faso (63%), as well as *A. arabiensis* (Burkina Faso, Cameroon, Tanzania) and *A. quadriannulatus* (Zambia). Resistance alleles occur as both homozygotes or heterozygotes in all species except *A. quadriannulatus*, which is always heterozygous (Figure 1B).

**Figure 1.**
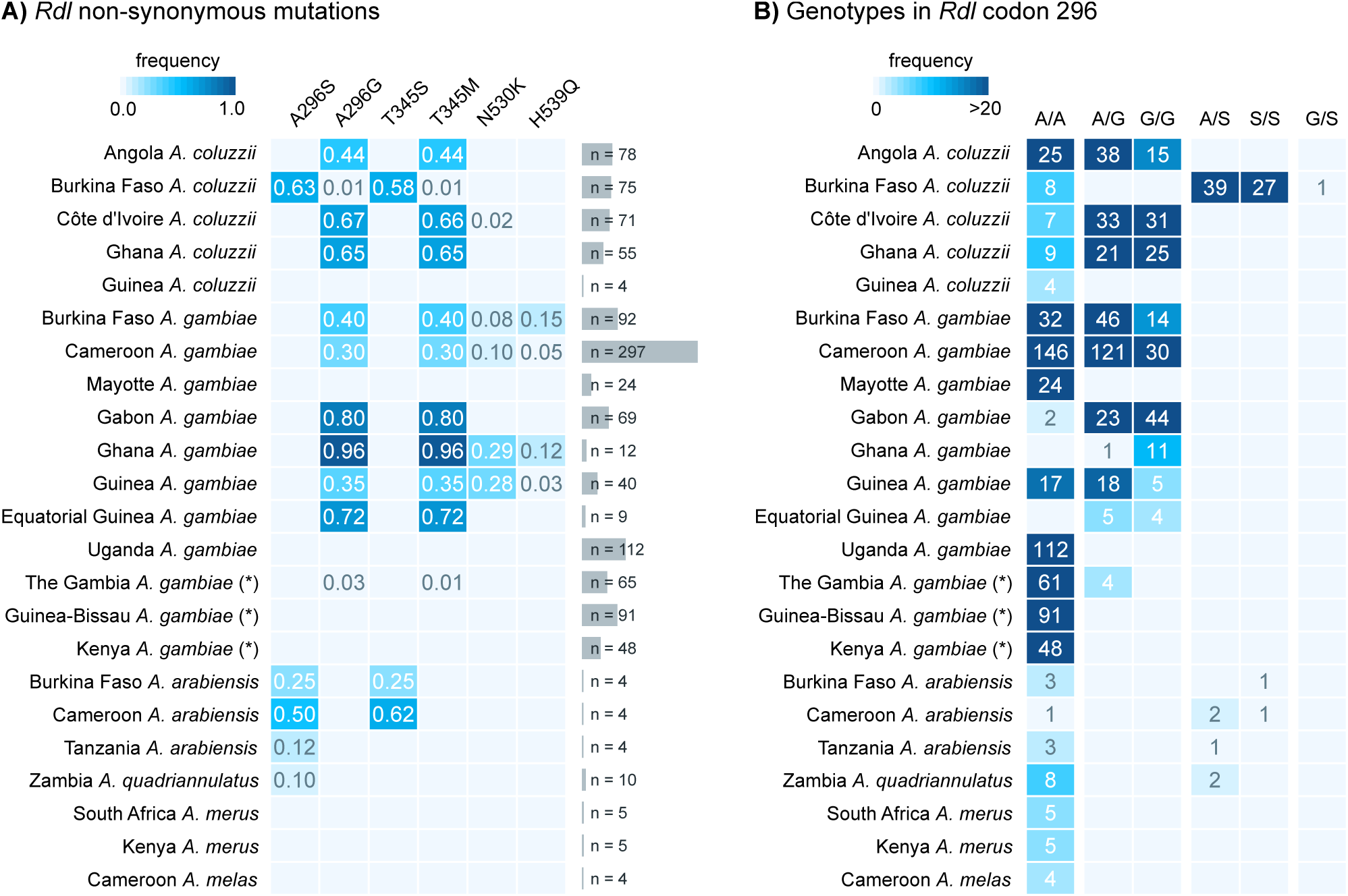
Rdl mutations. **A)** Frequency of non-synonymous mutations in *Rdl* across populations of *A. gambiae*, *A. coluzzii* (*Ag1000G* Phase 2) and *A. arabiensis*. Only variants with >5% frequency in at least on population are included. **B)** Distribution of genotypes for the two mutations in codon 296 (*A296S* and *A296G*). Note: *A. gambiae* populations denoted with an asterisk (The Gambia, Guinea-Bissau and Kenya) have high frequency of hybridisation and/or unclear species identification (see Methods).

We also identified two mutations in codon 345 with very similar frequencies to those of each codon 296 mutation: *T345M* (C-to-T in the second codon position), co-occurring with *A296G*; and *T345S* (A-to-T in the first codon position), co-occurring with *A296S*. The high degree of linkage disequilibrium between genotypes in codons 296 and 345 confirmed that they were co-occurring in the same specimens (Figure 2; e.g., the *296G*/*345M* allele pair had a Huff and Rogers *r* and Lewontin’s *D′* = 1), and was apparent in all individual populations where the alleles were present (Supplementary Material SM3). Codons 296 and 345 are located in the 7th and 8th exons of *Rdl*, separated by 3935 bp; and they map to the second and third transmembrane helices of the RDL protein, respectively (Supplementary Material SM4).

**Figure 2.**
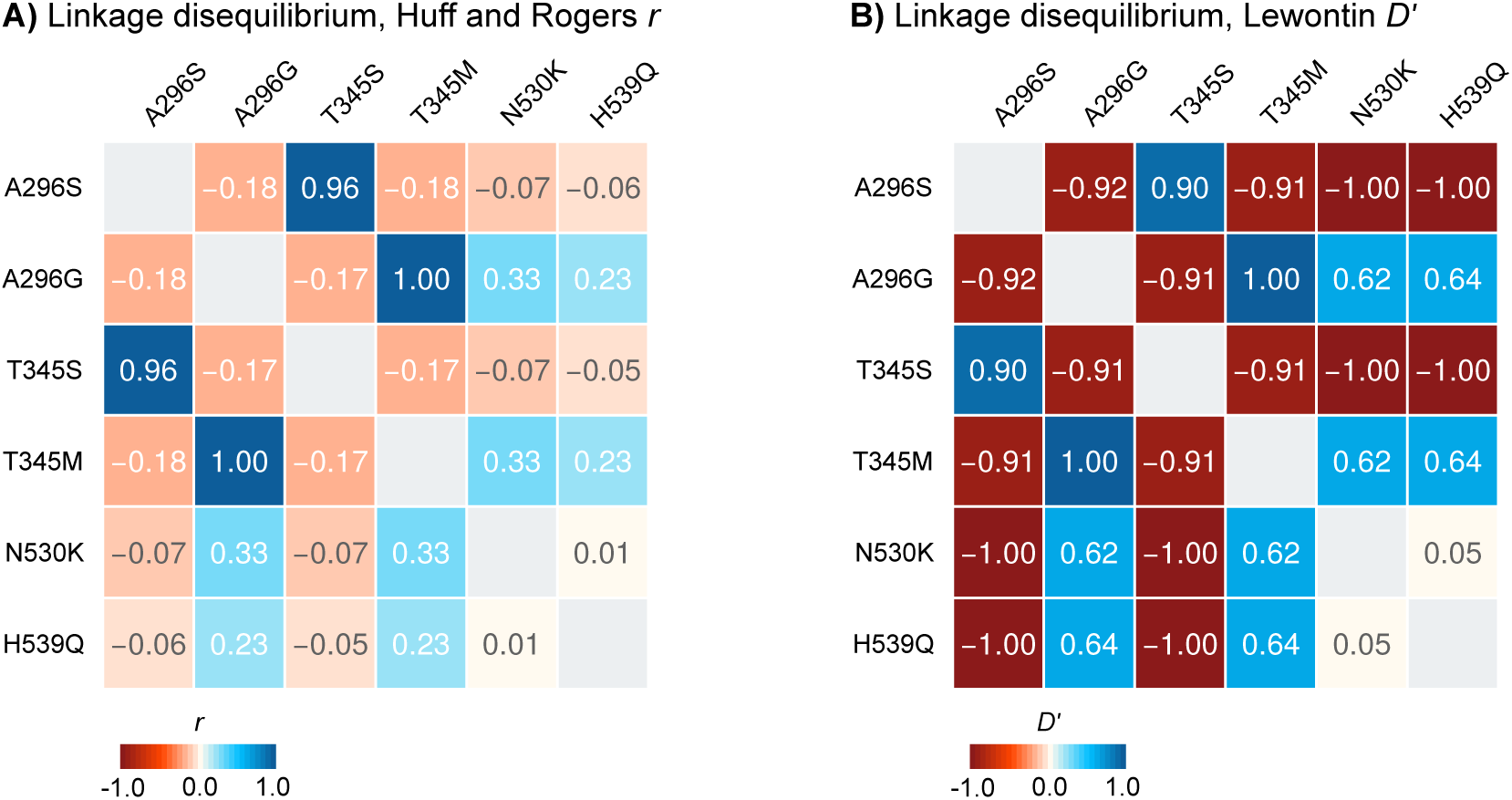
Linkage disequilibrium. Linkage disequilibrium between non-synonymous mutations in *Rdl*, calculated using Huff and Rogers’ *r* (A) and Lewontin’s *D′* (B).

### *Rdl* resistance mutations evolved on two unique haplotypes in *A. gambiae* and *A. coluzzii*

The high frequency of the *296S* and *296G* alleles in various populations of *A. gambiae* and *A. coluzzii* (Figure 1), together with their co-occurrence with nearby mutations (Figure 2), were suggestive of a selective sweep driven by positive selection on the resistance alleles. To clarify this possibility, we inspected the similarity of haplotypes in *A. gambiae*, *A. coluzzii* and the four outgroup species (*n* = 2356 haplotypes) using a minimum spanning network based on 626 phased variants located 10,000 bp upstream and downstream of codon 296 (Figure 3).

**Figure 3.**
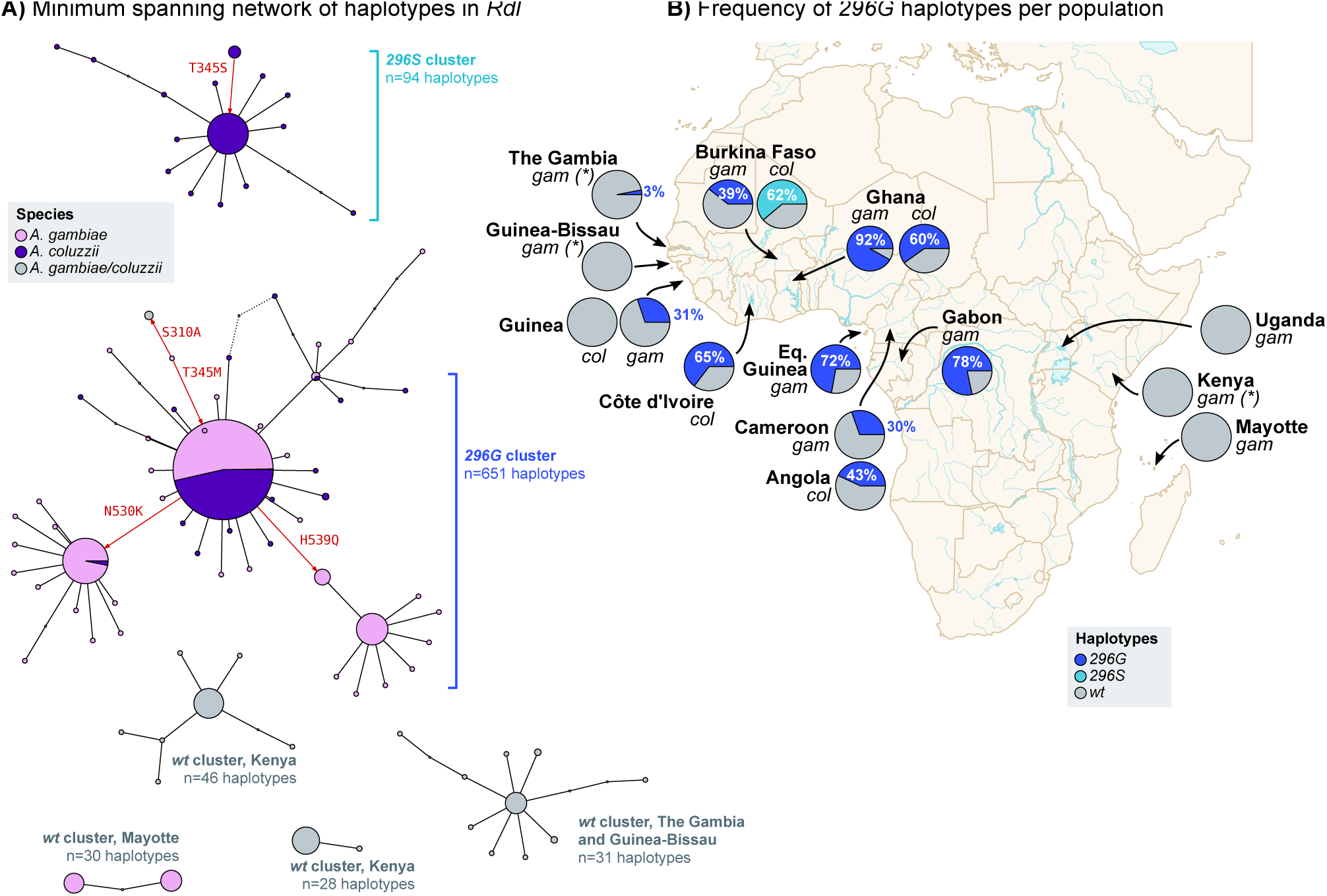
*Rdl* haplotypes. **A)** Minimum spanning network of haplotypes around *Rdl* codon 296 (626 phased variants located +/− 10,000 bp from the 2L:25429236 position). Only haplotype clusters with a frequency >1% in the cohort are represented (complete networks available as Supplementary Material SM6). Each node in the network is color-coded according to its species composition. Haplotype clusters carrying the resistance alleles *296G* and *296S* are highlighted in blue. Red arrows indicate the direction of non-synonymous mutations (relative to reference genome). **B)** Frequency of resistance haplotypes per population. Detailed frequencies with absolute counts in Supplementary Material SM14. Note: *gam*=*A. gambiae*, *col=A. coluzzii*; *gam* populations denoted with an asterisk have unclear species identification and/or high rates of hybridisation.

We identified two distinct groups of haplotypes associated with resistance mutations. First, the *296G* cluster contained haplotypes sharing the *296G*/*345M* alleles which were widely distributed in Central and West Africa (11 populations of *A. coluzzii* and *A. gambiae*; *n* = 651 haplotypes). The *296G* group showed two sub-clusters associated with the downstream mutations *N530K* and *H539Q* (red arrows in Figure 3A), which were present in a subset of mostly *A. gambiae* populations (Guinea, Ghana, Burkina Faso and Cameroon; Figure 1A); with just a few *A. coluzzii* from Côte d’Ivoire in the *N530K* cluster. Both *N530K* and *H539Q* are in partial linkage disequilibrium with *296G* alleles (Figure 2).

In contrast, the *296S* cluster, defined by ubiquitous co-occurrence of the *296S*/*345S* allele pair, was restricted to *A. coluzzii* from Burkina Faso (*n* = 94; Figure 3A, B), whereas the *A. arabiensis* and *A. quadriannulatus 296S* haplotypes appeared as distantly related singletons (not visible on Figure 3, see Supplementary Material SM5 and SM6). We also found four smaller wild-type clusters (*296A* allele; henceforth *wt*) that are specific to other geographical locations (Kenya, Mayotte, and The Gambia/Guinea-Bissau). The remaining haplotypes are also *wt* and group in smaller clusters or singletons with frequencies <1% in the dataset (*n* = 1476, 62.6% of all examined haplotypes; Supplementary Material SM5 and SM6).

Both the *296G* and *296S* haplotype clusters are often found in high frequencies within their respective populations. For example, *296S* was present in 62.3% of all Burkinabè *A. coluzzii*, and *296G* reached 91.7% in Ghanaian *A. gambiae* (Figure 3B).

The haplotype clustering analysis shows that all non-synonymous mutations (*T345M*, *T345S*, *N530K*, and *H539Q*) are associated with either the *296G* or the *296S* resistance haplotypes. The existence of seven non-synonymous mutations associated in haplotypes that have evolved over the last 70 years is remarkable: mosquito *Rdl* genes are highly conserved and have accumulated very few amino-acid mutations since anophelines diverged from culicines (for instance, *A. gambiae Rdl* retains a 97.6% amino-acidic identity with its *Aedes aegypti* ortholog and *d_N_/d_S_* = 0.052, indicating predominant purifying selection; Supplementary Material SM4). Here, we observe that the resistant haplotypes accumulate an excess of non-synonymous mutations compared to the *wt*, with non-synonymous to synonymous genetic diversity ratios (*π_N_/π_S_*) being ∼18x higher in the *296G* cluster (*π_N_/π_S_* = 2.428 +/− 0.009 standard error) than in *wt* haplotypes (*π_N_/π_S_* = 0.135 +/− 0.001); and ∼4x higher in *296S* (*π_N_/π_S_* = 0.485 +/− 0.018).

### The *296S* and *296G* alleles are associated with hard selective sweeps

Next, we investigated the signals of positive selection linked to the *296S* and *296G* resistance haplotypes. First, we found that haplotypes carrying *296G* and *296S* alleles had longer regions of high extended haplotype homozygosity (*EHH*) than the *wt* (Figure 4A), as expected under a scenario of selective sweeps linked to these resistant variants. A closer examination revealed that *EHH* decays slower at the 3′ region of *Rdl* (Figure 4A): in both clusters, *EHH* is above 0.95 (i.e. 95% of identical haplotypes) in the region downstream of codon 296 (exons 7 and 8), but decays more rapidly towards the 5′ of the gene (*EHH* < 0.20 in exon 6a/6b, *EHH* < 0.10 in exon 1). The core resistance haplotypes had lengths of 5,344 bp for *296G* and 4,161 bp for *296S* (defined at *EHH* > 95%), which were one order of magnitude higher than *wt* haplotypes (460 bp), and covered all non-synonymous mutations linked to codon 296 alleles (*T345M*, *T345S*, *N530K*, and *H539Q*).

**Figure 4.**
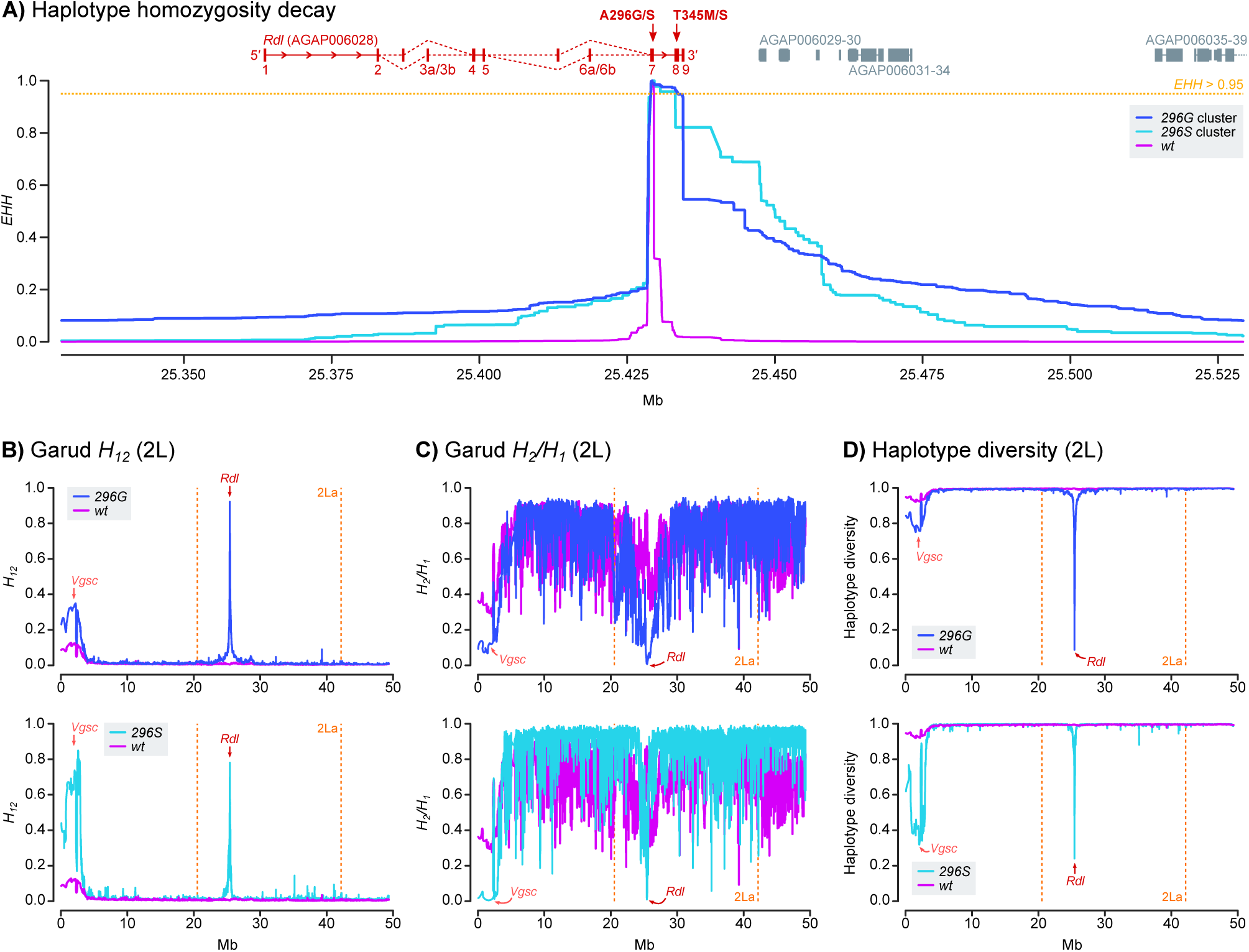
Positive selection of haplotypes carrying resistance mutations. **A)** Profile of *EHH* decay for each group of haplotypes (*296G*, *296S* and *wt*), built from 11,180 phased variants located +/− 100,000 bp from codon 296 (2L:25429236 position). Coordinates of nearby genes are indicated above the *EHH* panel (in *Rdl*, exons are numbered and red arrows indicate the position of codons 296and 345). **B-D)** Profiles of Garud *H_12_*, Garud *H_2_/H_1_* and haplotypic diversity along chromosomal arm 2L, highlighting the region covered by the 2La inversion (orange vertical lines) and the location of *Rdl* (red arrow). Each statistic was calculated separately for haplotypes carrying the *296G*, *296S* and *wt* alleles, using sliding blocks of 500 variants with 20% overlap.

Next, to estimate the softness/hardness of the sweep, we calculated the profile of Garud’s *H* statistics (Garud et al. 2015) and haplotypic diversity along the 2L chromosome arm (Figure 4B-D). Both *296G* and *296S* haplotype clusters showed signals of a hard selective sweep: (i) they had markedly higher Garud’s *H_12_* (*296G*: 0.698 +/− 0.001 standard error; *296S*: 0.744 +/− 0.006) than *wt* (0.003 +/− 0.0), which indicates an over-abundance of the most frequent haplotypes in the cohort (Messer and Petrov 2013; Garud et al. 2015); (ii) lower *H_2_/H_1_* ratios (*296G*: 0.052 +/− 0.0; *296S*: 0.011 +/− 0.007) than *wt* (0.756 +/− 0.001), indicative of a hard sweep with decreased background variation (Messer and Petrov 2013; Garud et al. 2015); and (iii) low haplotypic diversity (*296G*: 0.501 +/− 0.001; *296S*: 0.377 +/− 0.007) compared to the *wt* (0.998 +/− 0.000).

Unexpectedly, chromosomes containing *296G* and *296S* alleles also exhibited signals of positive selection at a distant pericentromeric region of 2L (Figure 4B-D), typically associated with strong selective sweeps around two mutations in the *Vgsc* gene (*995F* and *995S*) (Lynd et al. 2010; Clarkson et al. 2014; Clarkson et al. 2018), which is the target site of pyrethroids and DDT (Davies et al. 2007). Positive selection in *Vgsc* was particularly strong in chromosomes that also carried *296S* alleles (*H_12_* = 0.917 +/− 0.004 standard error), followed by *296G* (*H_12_* = 0.412 +/− 0.001) and, to a lesser degree, *wt* (*H_12_* = 0.147 +/− 0.000). However, neither of the *Vgsc* resistance alleles (*995F* and *995S*) are in linkage disequilibrium with *296G* or *296S* (Supplementary Material SM7, SM8).

Rather, this apparent association is due to geographical overlap: *296G* and *296S* are present in West African populations that are near-fixed for *Vgsc* resistance alleles (>80% *995F* in 7 out of 10 populations; Supplementary Material SM8), but are mostly absent elsewhere. Overall, *Rdl* resistance alleles are found on two unique sets of highly similar haplotypes (Figure 3), each of them specific to one allele (*296S* and *296G*), that underwent independent hard selective sweeps (Figure 4).

### Co-segregation of *Rdl* haplotypes and 2La inversions

*Rdl* lies within the 2La chromosomal inversion, which is the longest in the *A. gambiae* genome (20.5-42.1 Mb) (Coluzzi 2002). The 2La inversion emerged in the last common ancestor of the *A. gambiae* species complex (Fontaine et al. 2015) and is currently polymorphic in *A. gambiae* and *A. coluzzii* (Stump et al. 2007), where it is linked to a range of important phenotypes including adaptation to human environments (Coluzzi et al. 1979), aridity (Cheng et al. 2012), insecticide resistance (Weetman et al. 2018), and susceptibility to *Plasmodium falciparum* (Riehle et al. 2017). Given that recombination is strongly reduced between chromosomes with discordant inversion karyotypes (Andolfatto et al. 2001; Ayala and Coluzzi 2005; Kirkpatrick 2010), any assessment of the evolution of genes within the 2La inversion, such as *Rdl*, needs to take into consideration whether haplotypes reside in inverted (2La) or non-inverted (2L+^a^) backgrounds.

To address this issue, we estimated the 2La inversion karyotypes for the *Ag1000G* Phase 2 samples using a principal component analysis of allele presence/absence in the inverted region (using genomes with known inversion karyotypes as a reference; Figure 5A and Supplementary Material SM1 and SM9). The first principal component clearly discriminated between each of the inversion genotypes (non-inverted 2L+^a^/2L+^a^ homozygotes, inverted 2La/2La homozygotes, and 2La/2L+^a^ heterozygotes). We used this information to compare the frequencies of 2La karyotypes with *Rdl* codon 296 genotypes (Figure 5B), and the karyotype frequencies per population (Figure 5C). The pan-African *296G* allele is present in all inversion karyotypes, but is more common in non-inverted backgrounds (73% of *296G*/*296G* homozygotes have 2L+^a^/2L+^a^ karyotypes; Figure 5B), in both *A. gambiae* and *A. coluzzii* populations (Figure 5C). On the other hand, *296S* alleles from *A. arabiensis* and Burkinabè *A. coluzzii* occur exclusively within the 2La inversion (100% of *296S/296S* homozygotes are in 2La/2La karyotypes; Figure 5B).

**Figure 5.**
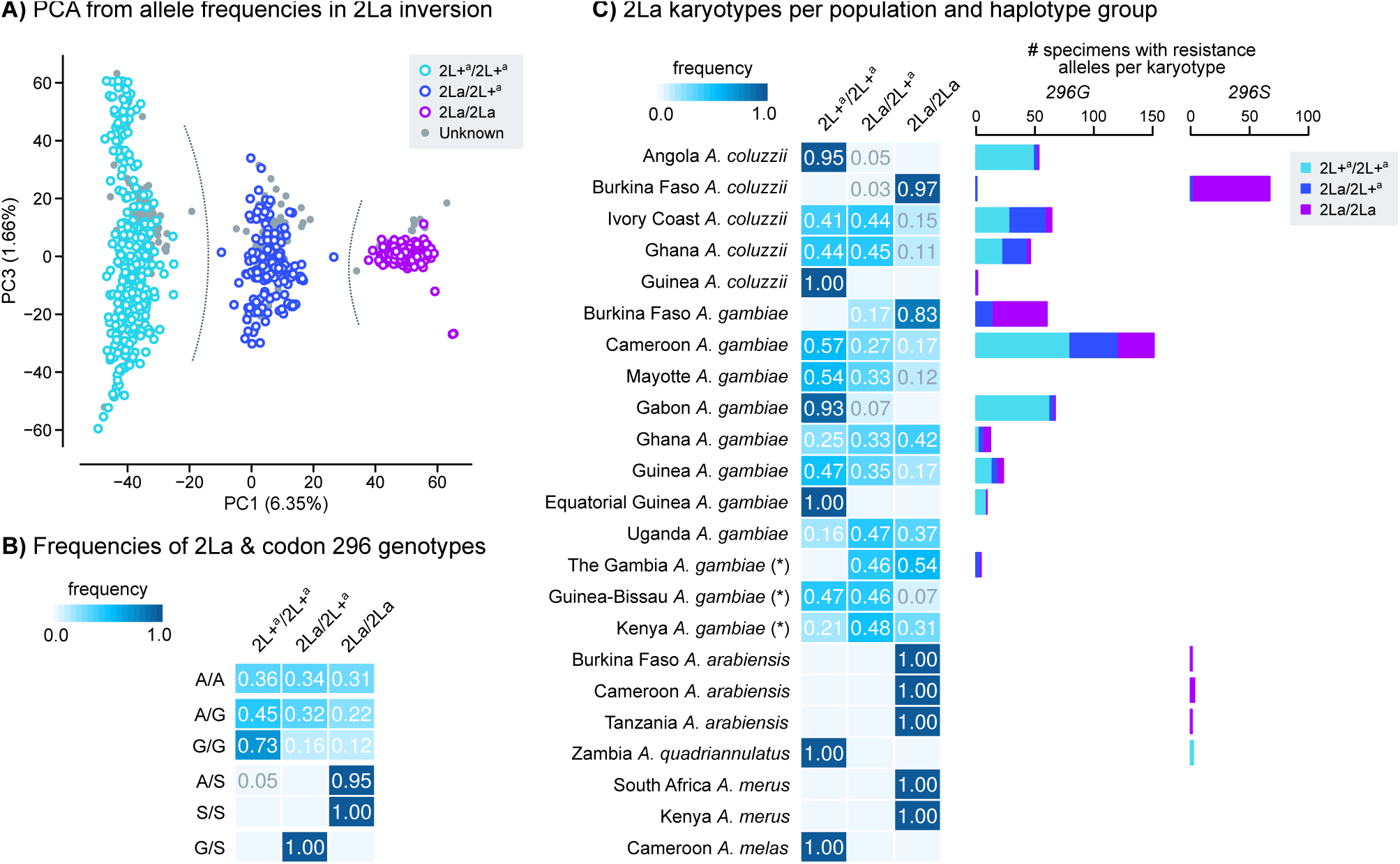
Genotypes of the 2La inversion. **A)** Principal component analysis of genotype frequencies of 10,000 random variants located within the 2La inversion (coordinates: 2L:20524058-42165532). Specimens from *Ag1000G* Phase 1 are color-coded by 2La karyotype (homozygotes and heterozygotes), and they are used as a reference to assign 2La genotypes to Phase 2 specimens (grey). Grey dotted lines highlight the separation of three clusters according to 2La karyotype. **B)** Frequency of 2La inversion and *Rdl* codon 296 genotypes. **C)** Frequency of 2La inversion karyotypes per population (heatmap, left), and number of specimens from each population carrying resistance alleles (*296G* and *296S*), broken down by 2La karyotype (barplots, right). Note: *A. gambiae* populations denoted with an asterisk (The Gambia, Guinea-Bissau and Kenya) have high frequency of hybridisation and/or unclear species identification (see Methods).

### Introgression of *Rdl* resistance haplotypes

In order to obtain a more complete picture of possible introgression events, we performed a phylogenetic analysis of haplotype alignments at four loci around *Rdl*: 5′ and 3′ regions of the gene, and two loci upstream and downstream of the gene body (Figure 6). These phylogenies highlight two events of interspecific introgression (explored below in grater detail): *296G* between *A. gambiae* and *A. coluzzii* (as reflected by their identical swept haplotypes; Figure 3), and *296S* between *A. coluzzii* and *A. arabiensis*. In addition, they also confirm the spread of *296G* haplotypes across different 2La inversion types (interkaryotypic introgression; Figure 5). In the following paragraphs, we characterise these introgressions and attempt to identify the donors and acceptors of each event.

**Figure 6.**
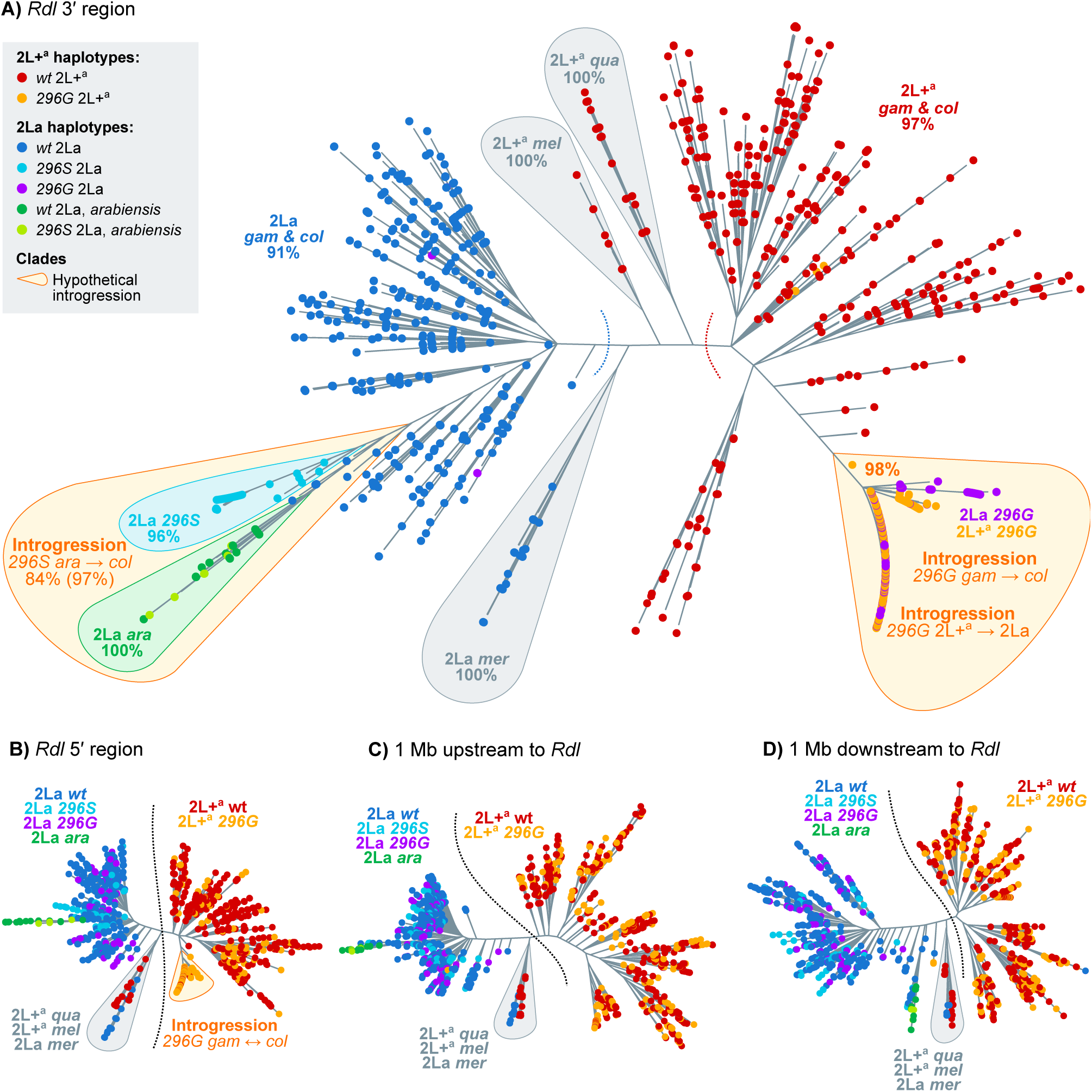
Phylogenies of haplotypes around the *Rdl* locus. **A)** Maximum-likelihood phylogenetic analysis of variants present at the 3′ region of *Rdl* (20,000 kbp). Nodes are haplotypes and have been color-coded according to their *Rdl* genotype (*296S*, *296G*, *wt*), 2La karyotype (2La, 2L+^a^) and species. Orange bubbles highlight clades with hypothetical introgression events. Grey bubbles highlight outgroup clades. Statistical supports are shown on selected clades (UF bootstrap). **C-E)** Analogous phylogenies from the *Rdl* 5′ region, upstream, and downstream regions within the 2La inversion (+/− 1 Mb of *Rdl*). Complete alignments and phylogenies in Supplementary Material SM10 and SM11. Species abbreviations: *col=coluzzii, gam=gambiae, ara=arabiensis, mer=merus; mel=melas, qua=quadriannulatus.* Arrows indicate introgression events.

### Interspecific introgression of *296G* and *296S* haplotypes

All four phylogenies exhibit two main clades separating *A. gambiae* and *A. coluzzii* haplotypes according to their 2La inversion karyotype, rather than by species (2La in blue, left; 2L+^a^ in red, right; ultrafast bootstrap support [UFBS] 91% and 97% respectively; Figure 6A). This clustering is due the fact that the 2La inversion has been segregating in *A. gambiae* and *A. coluzzii* since before the beginning of their speciation (Fontaine et al. 2015).

A closer examination shows that *Rdl*-specific phylogenies (Figure 6A, B) have a distinct sub-clade within the 2La cluster, consisting of *A. coluzzii 296S* haplotypes and *A. arabiensis*, some of which also possess the *296S* allele (light blue and green sequences in Figure in Figure 6A; UFBS 97%, 84% for their sister-branch relationship). The deep branching of *A. arabiensis* haplotypes within the *A. gambiae*/*coluzzii* 2La clade is to be expected, as *A. arabiensis* 2La inversions descend from an ancient introgression event from the *A. gambiae*/*coluzzii* ancestor (Fontaine et al. 2015). However, their close phylogenetic relationship with *A. coluzzii 296S* haplotypes is suggestive of interspecific introgression.

To confirm this event of introgression and ascertain its direction, we compared the results of two complementary Patterson’s *D* tests (Figure 7). The *D* statistic compares allele frequencies between three putatively admixing populations (A, B and C) and one outgroup (O), and can identify introgression between populations A and C (in which case *D* > 0) or B and C (*D* < 0; see Methods and (Durand et al. 2011; Patterson et al. 2012)).

**Figure 7.**
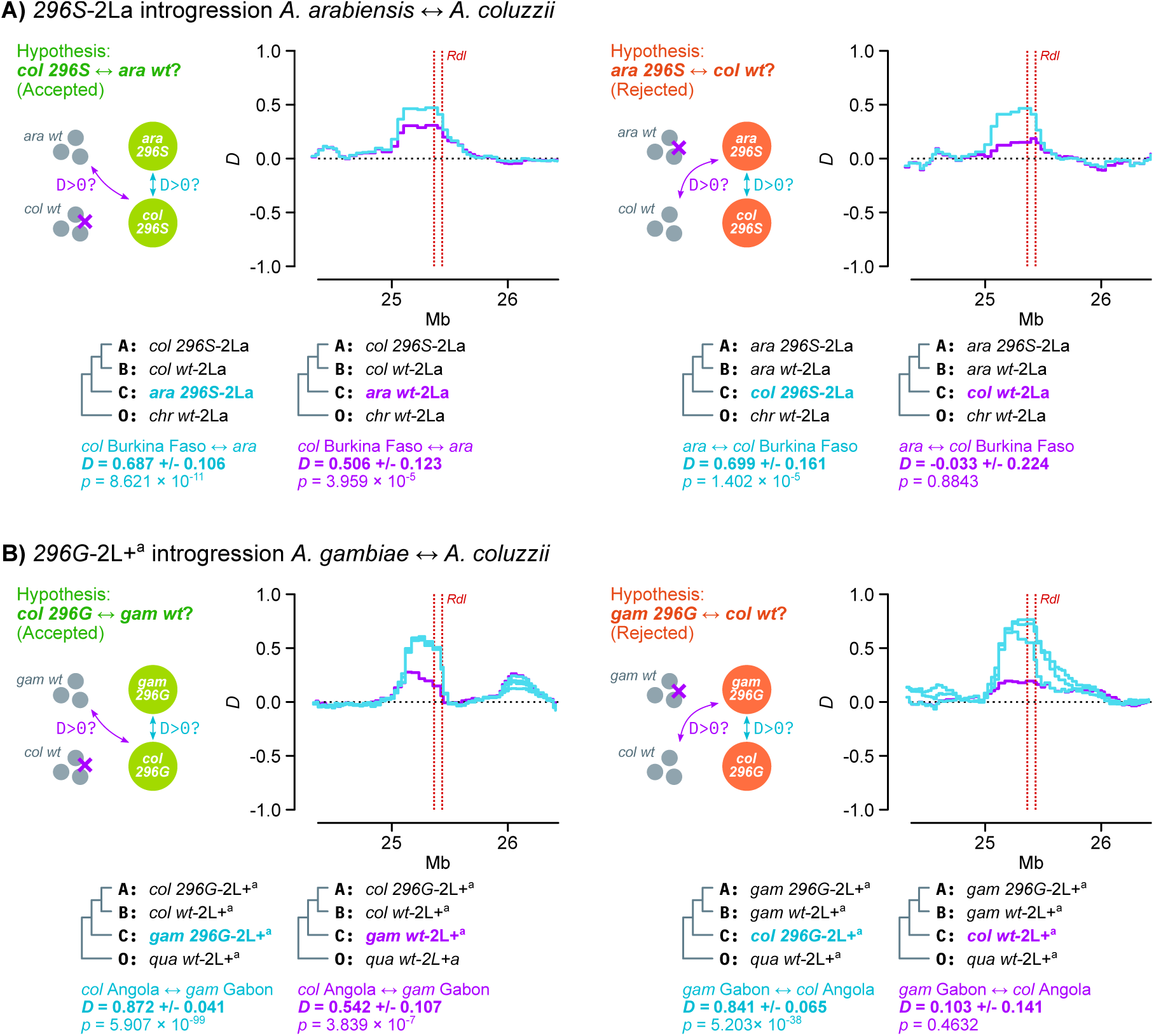
Interspecific introgression. **A)** Direction of *296S* introgression between *A. arabiensis* and *A. coluzzii* (2La/ 2La background). We test two complementary hypothesis using Patterson’s *D* statistics: left, introgression between *A. coluzzii 296S* homozygotes (population A), *A. coluzzii wt* (B) and *A. arabiensis* (*296S* or *wt*; C) using *A. christyi* as outgroup (O); right, reversing the position of *A. coluzzii* and *A. arabiensis* as populations A/B and C. The complementary hypotheses can be summarised as follows: if *296S* homozygotes from species *i* show evidence of introgression with *wt* homozygotes from species *j* (first test) but not with *wt* from species *i* (second test), *296S* originated in species *j*. **B)** Direction of *296G* introgression between *A. gambiae* and *A. coluzzii* (2L+^a^/2L+^a^ background), testing two complementary hypothesis using Patterson’s *D* statistics: left, introgression between *A. coluzzii 296G* homozygotes (population A), *A. coluzzii wt* (B) and *A. gambiae* (*296G* or *wt*; C) using *A. quadriannulatus* as outgroup (O); right, reversing the position of *A. coluzzii* and *A. gambiae* as populations A/B and C. Color-coded cladograms at the bottom of each plot indicate the groups of specimens used in each test, including the average *D* in the *Rdl* locus with standard errors and *p*-values (estimated from the Z-score of jack-knifed estimates; see Methods). See detailed lists of comparisons and statistical analyses in Supplementary Material SM12 and SM13.

Here, if *296S* had emerged in *A. arabiensis* and later introgressed into *A. coluzzii*, we would expect *296S A. coluzzii* specimens to exhibit *D* > 0 when compared to *296S A. arabiensis*, but also to be more similar to *wt A. arabiensis* (from which *296S* evolved) than to *wt A. coluzzii*. As predicted, we identify evidence of introgression between *A. coluzzii 296S* homozygotes and both (i) *296S A. arabiensis* (*D* = 0.687 +/− 0.106 standard error, *p* = 8.621 × 10^−11^ derived from a *Z*-score distribution) and (ii) *wt A. arabiensis* (*D* = 0.506 +/− 0.123, *p* = 3.959 × 10^−5^; left panel in Figure 7A). Conversely, if *296S* had introgressed from *A. coluzzii* into *A. arabiensis*, we would see evidence of introgression between *296S A. arabiensis* and *wt A. coluzzii*, but we do not (right panel in Figure 7A; *D* = −0.033 +/− 0.224, *p* = 0.884). These results are robust to various choices of outgroup species (*A. christyi* and *A. epiroticus*), and tests involving a negative control with fixed 2La inversions (*A. merus*) do not show evidence of introgression with *296S* specimens (Supplementary Material SM12). Thus, we conclude that the *296S* allele originated in *A. arabiensis* and later spread into *A. coluzzii*.

*Rdl* phylogenies (Figure 6A, B) also show a sub-clade of highly similar *A. gambiae* and *A. coluzzii* haplotypes within the 2L+^a^ cluster, all of them carrying *296G* alleles. This clade corresponds to the swept haplotypes identified above (Figure 3). We established the polarity of introgression using complementary Patterson’s *D* tests. Here, we found that *296G* haplotypes from resistant *A. coluzzii* populations (Côte d’Ivoire, Angola, and Ghana) exhibited signals of introgression with *wt A. gambiae* from Gabon (e.g. *D* = 0.542 +/− 0.107, *p* = 3.839 × 10^−7^ compared to Angolan *A. coluzzii*; Figure 7B); but that this signal of introgression disappeared when comparing *wt A. coluzzii* to *296G A. gambiae* from Gabon (e.g. *D* = 0.103 +/− 0.141, *p* = 0.4632 compared to Angolan *A. coluzzii*; Figure 7B) or elsewhere (Supplementary Material SM13). These results support the introgression of *296G* from *A. gambiae* to *A. coluzzii*.

The fact that only Gabonese *A. gambiae* have significant support as the *296G* donor population could indicate that they are closer to the founding *296G* haplotype and/or the original introgression event. However, the negative results in other populations harbouring *296G* alleles (Cameroon, Guinea; Supplementary Material SM13) could also be due to methodological limitations of our analysis – e.g., our conservative approach is restricted to specimens that are homozygous for both the inversion karyotype (2L+^a^/2L+^a^) and codon 296 (*296G*/*296G* or *wt*/*wt*); and the similarity between *wt A. gambiae* and *A. coluzzii* relative to the highly divergent swept haplotype can hinder the identification of the original background.

### The *296G* haplotype spread from 2L+^a^ to 2La chromosomes

The haplotype phylogeny from the *Rdl* 3′ region, where codon 296 variants reside, also revealed that the 2L+^a^ clade (non-inverted, red; Figure 6A) contained a sub-cluster of *296G* haplotypes from both 2L+^a^ (orange) and 2La orientations (purple; Figure 6A; UFBS 98%). The deep branching of *296G*-2La haplotypes within the 2L+^a^ clade implies that *296G* originated in a non-inverted background and later spread to inverted chromosomes via interkaryotypic introgression. Chromosomal inversions are strong barriers to recombination, but double cross-overs or gene conversion events can result in allelic exchange between non-concordant inversions (Andolfatto et al. 2001; Kirkpatrick 2010) and thus explain this phylogenetic arrangement.

However, the phylogeny of *Rdl* 5′ haplotypes (which excludes codon 296 and the adjacent non-synonymous mutations) showed that *296G-*2La sequences (purple) branched within the *wt*-2La clade instead (blue; Figure 6B). Thus, interkaryotypic introgression only affects the swept haplotype at the 3′ end of *Rdl* (Figures 3 and 4), whereas the 5′ region is closer to the *wt*. We can confirm whether the introgression is specific to the 3′ swept haplotype by examining the profile of sequence divergence along the *Rdl* gene locus (*Dxy;* Figure 8). We expect *296G* haplotypes to be more similar to *wt*-2L+^a^ than to *wt-*2La, given that the *296G* allele first evolved in a 2L+^a^ background (blue line, *Dxy* ratio > 1; Figure 8). In the case of *296G* alleles from 2La chromosomes, this expectation holds at the 3′ region of *Rdl* but not at 5′ nor outside of the gene, where allele frequencies are more similar to the *wt-*2La (purple line, *Dxy* ratio < 1; Figure 8).

**Figure 8.**
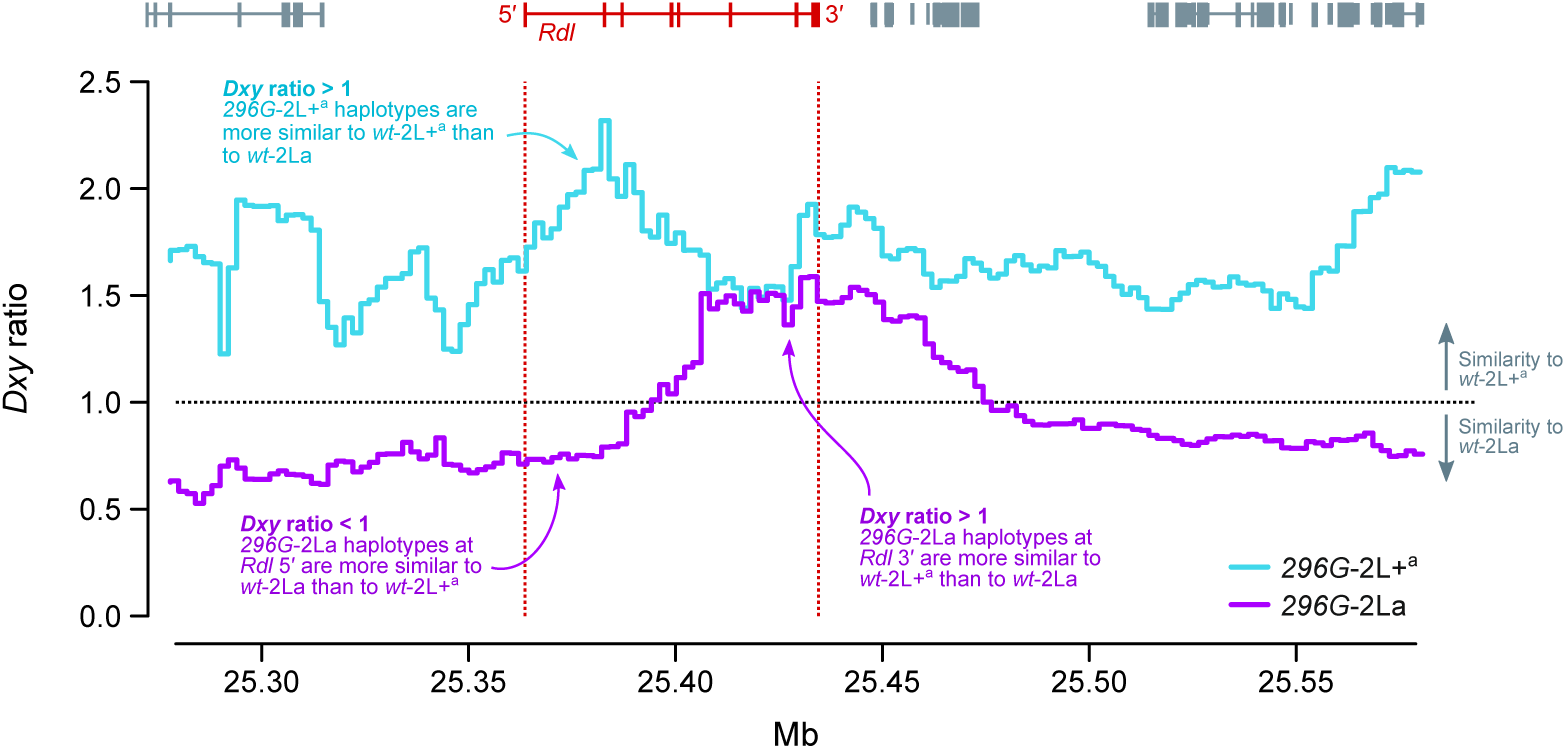
Interkaryotypic introgression of *296G* haplotypes. Ratio of sequence divergence (*Dxy*) between *296G* and *wt* haplotypes of 2L+^a^ and 2La origin. In this ratio, numerators are divergences between *296G* haplotypes (of either 2L+^a^ or 2La origin, in blue and purple respectively) relative to *wt-*2La haplotypes, and denominators are relative to *wt*-2L+^a^. Ratios >1 indicate similarity to *wt*-2L+^a^, and values <1 indicate similarity to *wt-2La.* All values are calculated in windows of 20,000 kbp with 10% overlap.

The presence of alleles from different karyotypic backgrounds in the *296G*-2La *Rdl* sequences is consistent with the sudden decay of haplotype homozygosity immediately upstream to codon 296 (Figure 4A), as the presence of *wt* alleles of 2La origin at 5′ of the *296G* swept haplotypes causes a faster decay in haplotype homozygosity in 2La than in 2L+^a^ haplotypes (Supplementary Material SM14A). Concordantly, haplotype diversity at the 5′ region of *Rdl* is higher in *296G*-2La than in *296G*-2L+^a^ haplotypes (Supplementary Material SM14B).

### Structural modelling predicts that *296G* and *296S* disrupt the dieldrin binding site in alternative ways

Finally, we investigated the effects of *296G* and *296S* resistance alleles on the structure of RDL receptors. The *A. gambiae* RDL receptor was modelled as a homopentamer based on the human GABA_A_ receptor structure (Masiulis et al. 2019) (Figure 9). In *wt* receptors, the *296A* residue is located near the cytoplasmic end of the pore-lining second transmembrane helix (M2) and its side chain is orientated into the pore (Figure 9A), whereas residue 345 is located distant from the pore, at the cytoplasmic end of the M3 helix with its side chain orientated towards the lipid bilayer. We carried out automated ligand docking for dieldrin in the *wt* receptor, finding a putative binding site along the receptor pore where the insecticide docked with estimated free energy of binding (*ΔG_b_*) of −8.7 kcal/mol (Figure 9B). The *296A* side chains form a major point of contact with the ligand. A structure of human GABA_A_ in complex with picrotoxin showed that this ligand forms multiple hydrogen bonds with residues lining the pore (Masiulis et al. 2019), but dieldrin lacks equivalent hydrogen bond-forming groups. Thus, the close contacts between *296A* side chains and dieldrin suggest that van der Waals interactions between these molecules are the predominant binding interaction.

**Figure 9.**
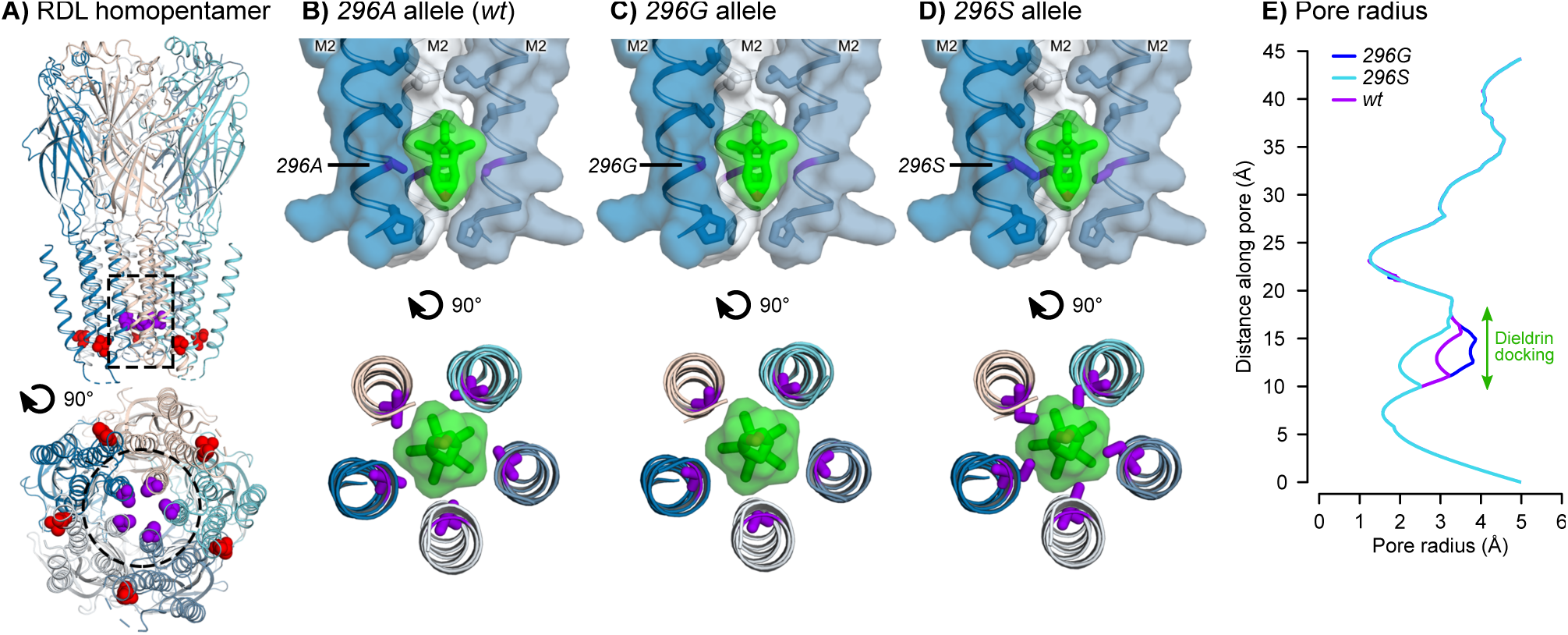
RDL receptor models with docked dieldrin. **A)** Homology model of the *A. gambiae* RDL homopentamer, viewed from the membrane plane (top) and cytoplasm (bottom). The *296A* (purple) and *345T* (red) positions are shown in space-fill. The dotted outlines depict the receptor regions in panels B-D. **B)** Docking prediction for dieldrin in the pore of the *296A* (*wt*) receptor. Dieldrin is shown in green, in sticks and transparent surface. Side chains lining the pore are shown as sticks and *296A* is coloured purple. **C-D)** Superimposition of dieldrin docking onto models of the *296G* and *296S* receptors, respectively. **E)** Pore radii in *296A*, *296G* and *296S* models.

Next, we superimposed the *wt* dieldrin docking coordinates onto models of resistant RDL receptors, resulting in disruptions of the predicted form of interaction (Figure 9C, D). The *A296G* substitution widens the pore at the dieldrin docking site (2.9Å to 3.8Å) and reduces the surface area of contact between the lumen and dieldrin (Figure 9C, E). *A296S* has the opposite effect: it results in a narrower pore (2Å) and shows an overlap between the serine side-chains and dieldrin, which indicates that steric hindrance could prevent the insecticide from binding at this location (Figure 9D, E).

## Discussion

### Evolution of *Rdl* resistance: selective sweeps and multiple introgression events

Contemporary dieldrin-resistant *A. gambiae* and *A. coluzzii* appear to descend from two unique hard selective sweeps around the *A296G* and *A296S* mutations (Figures 3 and 4). Both sweeps occurred independently on different genomic backgrounds (Figure 6), and have undergone at least three introgression events (Figures 6-8): (i) *296G* from *A. gambiae* to *A. coluzzii*; (ii) *296G* from 2L+^a^ to 2La chromosomes; and (iii) *296S* from *A. arabiensis* to *A. coluzzii*.

In the case of *296G*, our data supports an origin in *A. gambiae* with 2L+^a^ chromosomes, followed by interspecific introgression into *A. coluzzii*, and interkaryotypic introgression into 2La chromosomes. The *A. gambiae* origin is inferred from the background similarity between *A. coluzzii* swept haplotypes and *A. gambiae wt* specimens from Gabon (according to Patterson’s *D* test; Figure 7B). *A. gambiae* resistance haplotypes have accrued more non-synonymous mutations than *A. coluzzii* (*N530K* and *H539Q*; Figure 1A), which is consistent with a longer evolutionary history in the former. In either case, the swept haplotype currently spans populations of both species across West and Central Africa – mimicking the pan-African selective sweep described for the homologous *Rdl* mutation in *D. melanogaster* (ffrench-Constant, Rocheleau, et al. 1993; ffrench-Constant, Steichen, et al. 1993; Thompson et al. 1993). This result is in line with previous studies that had hypothesized the existence of a pan-African *296G* sweep due to the strong genetic differentiation found in this locus (Lawniczak et al. 2010).

The interkaryotypic introgression of *296G* haplotypes from non-inverted 2L+^a^ into 2La chromosomes (Figures 6 and 7) also facilitated the spread of *296G* resistance alleles, e.g. in *A. gambiae* populations with high frequencies of 2La/2La karyotypes such as Burkina Faso (Figure 5C). While it is generally acknowledged that chromosomal inversions strongly suppress recombination (Sturtevant 1917), genetic exchange can occur via double cross-over recombination or gene conversion (Chovnick 1973; Rozas and Aguadé 1994; Andolfatto et al. 2001; Kirkpatrick 2010). The reduction in recombination is weaker in regions distant from the inversion breakpoints (Andolfatto et al. 2001), as it is the case for *Rdl* (located ∼4.8 Mb and ∼16.7 Mb away from the 2La breakpoints), which results in reduced differentiation at the centre of the inversion (Stump et al. 2007; Cheng et al. 2012) (Supplementary Material SM15). To the best of our knowledge, reports of adaptive introgression of individual genes within inversions are rare. In *Anopheles*, one of such cases are certain loci involved in adaptation to desiccation, which are linked to 2La inversions but are exchanged in 2La/2L+^a^ heterozygotes (Cheng et al. 2012; Ayala et al. 2019). Another example, possibly linked to gene conversion, could be the *APL1* cluster of hyper-variable immune genes: their pattern of sequence variation is more strongly influenced by geography and species (*A. gambiae*/*A. coluzzii*) than by the 2La inversion where they reside (Rottschaefer et al. 2011).

On the other hand, the *296S* selective sweep has a more restricted geographical distribution. In the *Ag1000G* cohort, *296S* is only found in *A. coluzzii* from Burkina Faso (Figure 3). We also identify *296S* alleles in *A. arabiensis* specimens from East (Tanzania), Central (Cameroon) and West Africa (Burkina Faso); as well as two *A. quadriannulatus* specimens from Zambia (which appears to be the first record in this species; Figure 1B).

Interestingly, we find clear evidence of *296S* introgression from *A. arabiensis* into *A. coluzzii* even when comparing to *A. arabiensis wt* specimens (Figure 7A), and despite the fact that none of the *A. arabiensis 296S* share the *A. coluzzii* swept haplotype (Figures 3A, 6A, and Supplementary Material SM6). Thus, lack of genomic evidence from *A. arabiensis* precludes the identification of the actual donor haplotype. A wider sampling of *A. arabiensis* populations will be necessary to complete the picture of *296S* evolution, in order to (i) identify the number of historical *A296S* mutations in this species; (ii) establish whether they were associated with one or more selective sweeps; and (iii) whether any of these hypothetical sweeps introgressed into *A. coluzzii*.

### Persistence of *Rdl* mutations after dieldrin withdrawal

Rdl is a highly conserved gene, with an extreme paucity of non-synonymous mutations over >100 Mya of evolutionary divergence (Neafsey et al. 2015) in culicines and anophelines, and low d_N_/d_S_ ratios that indicate a prevalence of purifying selection (Supplementary Material SM4). In this context, the persistence of 296G and 296S alleles in natural populations for more than 70 years, in spite of its fitness costs in the absence of insecticide (Rowland 1991a; Rowland 1991b; Platt et al. 2015), has been a long-standing puzzle.

Our study provides two key insights to this question. First, we find that, relative to the wt, haplotypes with resistance alleles have an excess of non-synonymous genetic diversity (∼18x increase in π_N_/π_S_ in 296G, ∼4x in 296S). This observation suggests that the emergence of 296G and, to a lesser degree, 296S, has substantially altered the selective regime of Rdl and enabled the accumulation of additional non-synonymous mutations in an otherwise highly constrained protein. A similar change has been recently observed for kdr mutations in Vgsc (the target site of pyrethroids), whereby 995F resistance haplotypes accumulate an excess of amino-acidic substitutions (Clarkson et al. 2018).

Second, we identify a high degree of genetic linkage between the *296G*/*345M* and *296S*/*345S* allele pairs, which is observed in all West African populations where codon 296 mutations are present (Figures 1, 2, and Supplementary Material SM3) due to the fact that virtually all swept haplotypes include both mutations (Figures 3 and 4). This near-universal association is highly relevant because codon 345 mutations are suspected to have compensatory effects that offset the costs of codon 296 variants (Remnant et al. 2014; Taylor-Wells et al. 2015). Studies of fipronil resistance have shown that both the *296G* allele and the combination of *296G* and *345M* alleles resulted in decreased insecticide sensitivity in *A. gambiae* (Taylor-Wells et al. 2015), *D. melanogaster* (Remnant et al. 2014), and *D. simulans* (Le Goff et al. 2005). Crucially, Taylor-Wells *et al*. (Taylor-Wells et al. 2015) showed that, in addition to fipronil resistance, the *A. gambiae 296G* allele causes heightened sensitivity to the GABA neurotransmitter (possibly contributing to the observed fitness costs (Rowland 1991b; Rowland 1991a; Platt et al. 2015)); and that the addition of the *345M* mutation reduces these detrimental effects while still conferring resistance.

Interestingly, our structural modelling analyses predict opposite resistance mechanisms for each resistance allele: *296G* results in a wider RDL pore with reduced van der Waals interactions with dieldrin (Figure 9C, E); whereas *296S* narrows the pore and impedes dieldrin docking due to steric hindrance (Figure 9D, E). These two effects suggest the possibility that the mechanisms behind the hypothesised compensatory roles of codon 345 mutations could be different as well, and open a new line of inquiry to investigate the exclusive association of each resistance variant with downstream mutations (*296G* with *345M*, *296S* with *345S*). Yet, the exact nature of the interaction between these codon 296 and 345 mutations remains unclear. Firstly, residue 345 does not have direct contacts with dieldrin or residue 296 (Figure 9A). Secondly, indirect effects are uncertain too: in human receptors, mutations at the interface between the third and second transmembrane helices (where residues 345 and 296 reside, respectively) affect the transition to the desensitized functional state (Gielen et al. 2015); but residue 345 in *A. gambiae* is not buried in this interface and is instead facing the lipid bilayer (Figure 9A), and the predicted effects of mutations *T345M* and *T345S* are not obvious.

Other possible factors behind the persistence of *Rdl* resistance alleles include the long half-life of dieldrin as an environmental organic pollutant; as well as the fact that *Rdl* is the target site of other insecticides such as fipronil, isoxazoline or meta-diamides (Gant et al. 1998; Nakao and Banba 2015; Miglianico et al. 2018); a secondary target of avermectin (Miglianico et al. 2018), and, possibly, of neonicotinoids (imidacloprid), pyrethroids (deltamethrin) (Taylor-Wells et al. 2015), and DDT (Lucas et al. 2019).

### Implications for vector control

The apparent ease with which *Rdl* adaptive haplotypes have spread across the barriers to recombination posed by species isolation (*A. gambiae*/*A. coluzzii* and *A. arabiensis*/*A. coluzzii*) and non-concordant chromosomal inversions (2L+^a^/2La) mirrors previous findings in *Vgsc* target site mutations (Clarkson et al. 2014), and suggests worrying consequences for insecticide deployment programmes. Burkina Faso, where resistance alleles have traversed both barriers to recombination, is a case-in-point example of this risk: the high frequency of 2La inversions (Figure 5C) did not prevent the spread of *296G*, and interspecific introgression of *296S* from *A. arabiensis* compounded this problem in *A. coluzzii*. In the future, a similar scenario could facilitate the spread of *296S* in East African *A. gambiae* and *A. coluzzii*, via adaptive introgression from *A. arabiensis*.

Also noteworthy is the overlap of *Rdl* and *Vgsc* resistance variants in West and Central Africa. The lack of genetic linkage between *Vgsc* and *Rdl* resistance haplotypes suggests that this co-occurrence is purely geographical, and does not fit a hypothetical epistatic relationship (Supplementary Material SM7 and SM8). Yet, this overlap is still relevant for vector control: as pyrethroid resistance increases in *Anopheles* populations (Ranson et al. 2011), the search for substitutes should take into account that some can be rendered ineffective by *296S* or *296G* (e.g. fipronil (Gant et al. 1998), avermectin (Miglianico et al. 2018), or, possibly, neonicotinoids such as imidacloprid (Taylor-Wells et al. 2015)). This risk is currently highest in the West and Central African populations of *A. gambiae* and *A. coluzzii* where both *296G* and *Vgsc 995F* (Clarkson et al. 2018) are common (Supplementary Material SM8). In the future, the introgression of *296S* from East African *A. arabiensis* could further compound current complications caused by the already high frequencies of *Vgsc 995S* in this region (Clarkson et al. 2018).

This case study of the mechanisms that underlie persistence of dieldrin resistance is also relevant for integrated resistance management. Strategies such as insecticide rotations or mosaics rely on a gradual decline in resistance over time (World Health Organization 2012). Instead, *296G* and *296S* haplotypes have accumulated additional non-synonymous mutations (Figure 3A), some of which (codon 345) are putatively compensatory. As mentioned above, a similar altered selective regime has also been observed in *Vgsc* haplotypes with *kdr* mutations (Clarkson et al. 2018). Interestingly, a study of Brazilian *Aedes aegypti* found that *Vgsc kdr* mutations did not decrease in frequency after a decade without public pyrethroid spraying campaigns (Macoris et al. 2018).

Brazilian *Aedes* have a longer history of pyrethroid-based treatments than African *Anopheles spp.* (van den Berg et al. 2012; Macoris et al. 2018); thus, their resilient *kdr* mutations could be (i) recapitulating our observations with respect to *Rdl* and dieldrin, and (ii) prefiguring a similar persistence of *Vgsc kdr* in the *A. gambiae* complex after a future phasing-out of pyrethroids in response to their decreasing efficacy (Ranson et al. 2011).

Overall, our results show that the *Rdl* resistance mutations that appeared after the pioneering deployment of dieldrin in the 1950s will still be relevant in the immediate future. Continued monitoring is thus necessary to understand the evolving landscape of genomic variation that underlines new and old mechanisms of insecticide resistance.

## Methods

### Data collection

We used variation data from individual *A. coluzzii* and *A. gambiae* mosquitoes from the *Anopheles gambiae* 1000 Genomes online archives, for Phase 2-AR1 (The *Anopheles gambiae* 1000 Genomes Consortium 2017). Specifically, we retrieved the phased genotype calls, SNP effect predictions, and the array of accessible genomic positions. We also obtained the same data for populations of four species in the *Anopheles* complex (*A. arabiensis*, *A. quadriannulatus*, *A. melas* and *A. merus*) and two outgroups (*A. epiroticus* and *A. christyi*), as available in the *Ag1000G* online archive (The *Anopheles gambiae* 1000 Genomes Consortium 2017). The complete list of downloaded genomes with accession codes is available in Supplementary Material SM1.

The reference gene annotation of *A. gambiae* was obtained from Vectorbase (Giraldo-Calderón et al. 2015) (GFF format, version AgamP4.9). Gene and variant coordinates employed in this study are based on the AgamP4 version of the genome assembly.

### Genotype frequencies and linkage disequilibrium

We retrieved all non-synonymous genomic variants located within the coding region of *Rdl* (genomic coordinates: 2L:25363652-25434556) that were biallelic, phased, and segregating at >5% frequency in at least one population (henceforth, ‘non-synonymous variants’). Parsing and filtering of genotype calls from *Ag1000G* was done using the *scikit-allel* 1.2.1 library (Miles and Harding 2017) in Python 3.7.4.

We calculated the linkage disequilibrium between each pair of non-synonymous variants using (i) Rogers’ and Huff *r* correlation statistic (Rogers and Huff 2009), as implemented in *scikit-allel* (*rogers_huff_r*); and (ii) Lewontin’s *D′* statistic (Lewontin 1964), as implemented in (Clarkson et al. 2018).

### Haplotype networks

We constructed a network of haplotype similarity using 626 biallelic, phased and non-singleton (shared between more than two samples) variants located in a region +/− 10kbp of *Rdl* codon 296 (middle nucleotide, coordinate 2L:25429236). We used the presence/absence of each allele within this genomic region to calculate Hamming distances and build minimum spanning networks (Bandelt et al. 1999), using the *hapclust* function from (Clarkson et al. 2018) (with distance breaks >3 variants). Network visualizations were produced using the *graphviz* 2.38.0 Python library (Ellson et al.), with haplotype clusters being color-coded according to species, population and presence/absence of the resistance alleles in codon 296 (*296S,* 2L:25429235; *296G*, 2L:25429236) and the 995th codon of *Vgsc* (Figure 3, Supplementary Material SM5, and SM6). The network visualization in Figure 3A excludes singletons and haplotype clusters with a cohort frequency <1%.

We calculated the sequence diversity (*π*) of each haplotype group in the same region (*sequence_diversity* function in *scikit-allel*), using a jack-knife procedure (iterative removal of individual haplotypes without replacement) (Tukey 1958) to estimate the average and standard error. We also calculated the sequence diversity in non-synonymous coding variants from this region (*π_N_*), synonymous coding variants (*π_S_*), and their ratio (*π_N_/π_S_*).

### Positive selection in haplotype clusters

We analysed the signals of positive selection in three haplotype groups, divided according to alleles in codon 296: *wt* (*n* = 1476), *296S* (*n* = 94) and *296G* (*n* = 651) (Supplementary Material SM5). First, we calculated the extended haplotype homozygosity decay (*EHH*) of each group of haplotypes, using 22,910 variants (phased and biallelic) located +/− 200 kbp of codon 296 (2L:25429236) (using the *ehh_decay* utility in *scikit-allel*). For each haplotype group, we recorded the genomic region where *EHH* decay >0.95 and <0.05.

Second, we calculated the profile of Garud’s *H* statistics (Garud et al. 2015) along the 2L chromosomal arm (*moving_garud_h* utility in *scikit-allel*; block length = 500 phased variants with 20% step). We performed the same calculations for the haplotypic diversity (*moving_haplotype_diversity* in *scikit-allel*). We calculated the Garud *H* and haplotypic diversity estimates in the *Rdl* locus, using a jack-knife procedure (Tukey 1958) (iterative removal of individual haplotypes without replacement) to calculate the mean and standard error of each statistic.

### Karyotyping of 2La inversions

In order to assign karyotypes of the 2La inversion in all specimens from *Ag1000G* Phase 2, we used known 2La karyotypes from Phase 1 as a reference (Miles et al. 2017), and analysed genotype frequencies within the inversion by principal component analysis (PCA). Specifically, we retrieved the genotype frequencies of 1142 specimens from *Ag1000G* Phase 2, 765 of which were also present in Phase 1 and had been previously karyotyped for this inversion (Miles et al. 2017); and selected 10,000 random SNPs (biallelic, shared between more than two samples, phased, segregating in at least one population, and located within the 2La inversion 2L:20524058-42165532). SNPs fitting these criteria were selected using the *scikit-allel* Python library, and the PCA was performed using the *randomized_pca* utility (with Patterson scaling).

Manual inspection of the principal components (Supplementary Material SM9) showed that PC1 (6.35% of variance explained) was sufficient to discriminate between known karyotypes from Phase 1 using a clear-cut threshold (2La/2La, 2La/2L+^a^ and 2L+^a^/2L+^a^). We determined the optimal classification thresholds using the C-Support Vector classification method (SVC, a method for supervised learning) implemented in the *scikit-learn* 0.21.3 Python library (Pedregosa et al. 2011). Specifically, we used the *SVC* function in *scikit-learn* (*svm* submodule) to train a classifier with known karyotypes from Phase 1 (765 observations) and the main principal components of the PCA analysis (10 variables), using a linear kernel and C=1. The selected thresholds were able to classify Phase 1 data into each of the three categories (2La/2La, 2La/2L+^a^ and 2L+^a^/2L+^a^) with 100% accuracy (as per the classifier *score* value), precision and recall (calculated using the *classification_report* function from the *scikit-learn metrics* submodule).

### Phylogenetic analysis of haplotypes

We obtained genomic alignments of SNPs located from four regions around the *Rdl* locus, at the following coordinates: (i) 5′ start of the gene (2L:25363652 +/− 10,000 kbp, 696 variants), (ii) 3′ end of the gene (2L:25434556 +/− 10,000 kbp, 428 variants), (iii) unadmixed region 1Mb upstream of *Rdl* (2L:24363652 + 20,000 kbp; 2903 variants; inside of the 2La inversion), and (iv) unadmixed region 1Mb downstream of *Rdl* (2L:26434556 + 20,000 kbp, 2594 variants; inside of the 2La inversion). These alignments were built from phased, biallelic variants within the aforementioned regions, obtained from *A. coluzzii* and *A. gambiae* (*Ag1000G* Phase 2), *A. arabiensis*, *A. quadriannulatus*, *A. melas* and *A. merus*. We restricted our analysis to haplotypes pertaining to individuals homozygous for the 2La inversion (2La/2La and 2L+^a^/2L+^a^), totalling 1684 haplotypes (out of 2356 haplotypes in the original dataset, obtained from 1178 specimens). Invariant sites were removed from the alignments using *snp-sites* 2.3.3 (Page et al. 2016). All alignments are available in Supplementary Material SM10.

Each genomic alignment was then used to compute Maximum-Likelihood phylogenetic trees using *IQ-TREE* 1.6.10 (Nguyen et al. 2015). The best-fitting nucleotide substitution model for each alignment was selected using the *TEST* option of *IQ-TREE* and according to the Bayesian Information Criterion (BIC), which suggested the GTR substitution matrix with ascertainment bias correction, four gamma (*Γ*) rate categories, and empirical state frequencies observed from the alignment (F) (i.e. the *GTR+F+ASC+G4* model in *IQ-TREE*). We calculated branch statistical supports using the UF bootstrap procedure (Minh et al. 2013; Hoang et al. 2018) and refined the tree for up to 10,000 iterations, until convergence was achieved (correlation coefficient ≥ 0.99).

Tree visualizations were created in R, using the *plot.phylo* function from the *ape* 5.3 library (Paradis and Schliep 2019) and *stringr* 1.4.0 (Wickham 2019). Each phylogeny was midpoint-rooted with *phytools* 0.6-60 (Revell 2012) (*midpoint.root*), and branch lengths in Figure 6 were constrained for display purposes (5 × 10^−5^ to 5 × 10^−3^ per-base substitutions range; unmodified trees available in Supplementary Material SM11).

### Interspecific introgression with Patterson’s *D* statistic

We analysed the signals of introgression along the 2L chromosomal arm using Patterson’s *D* statistic (Durand et al. 2011; Patterson et al. 2012). This statistic requires allele frequencies in four populations (A, B, C and O) following a predefined (((A,B),C),O) phylogeny, where A, B and C are populations with possible introgression events, and O is an unadmixed outgroup. Then, *D* > 0 if there is an excess of allele frequency similarities between A and C (which means either A → C or C → A introgression) and *D* < 0 for excess of similarity between B and C (B → C or C → B introgression) (Durand et al. 2011; Patterson et al. 2012). We calculated Patterson’s *D* along blocks of adjacent variants in the 2L chromosomal arm (block length = 10,000 variants, with 20% step length; phased variants only) using the *moving_patterson_d* utility in *scikit-allel*. We also calculated *D* in the *Rdl* locus (2L:25363652-25434556), and estimated its deviation from the null expectation (no introgression: *D* = 0) with a block-jackknife procedure (block length = 100 variants; *average_patterson_d* in *scikit*-*allel*). We then used these jack-knifed estimates to calculate the standard error, *Z*-score and the corresponding *p* value from the two-sided *Z*-score distribution.

Using the procedure described above, we performed multiple analyses of introgression between combinations of populations fitting the (((A,B),C),O) phylogeny. For each analysis, we selected A, B, C and O populations according to two criteria: (i) which interspecific introgression event was under test (*A. gambiae ∼ A. coluzzii* or *A. coluzzii ∼ A. arabiensis*); (ii) homozygous karyotypes of the 2La inversion within which *Rdl* is located (given that it introduces a strong effect on genotype frequencies across the entire *A. gambiae* species complex (Fontaine et al. 2015)) and the resistance haplotype in question; and (iii) exclude populations with high frequencies of hybrids, with controversial species identification, or with extreme demographic histories (Guinea-Bissau, The Gambia, and Kenya) (Miles et al. 2017; Vicente et al. 2017). Following these criteria, we then tested the presence and direction introgression between the combinations of populations specified below.

First, we tested the *A. coluzzii* ∼ *A. arabiensis* introgression of the *296S* haplotype in inverted genomes (2La/2La homozygotes; Figure 7A and Supplementary Material SM12). We performed two versions of this test, using either *A. coluzzii* or *A. arabiensis* as donors (population C), which can give an indication of the population of origin of the *296S* mutation. First, we tested the *A. arabiensis* → *A. coluzzii* hypothesis using: (i) *296S* homozygous *A. coluzzii* from Burkina Faso as population A; *wt* homozygous *A. coluzzii* from Burkina Faso as population B; (iii) *A. arabiensis* and *A. merus* specimens as multiple C populations (donors) C, treating *296S* and *wt* homozygous specimens as different populations; and (iv) *A. epiroticus* and *A. christyi* as population O. Second, we tested the *A. coluzzii* → *A. arabiensis* hypothesis but switching the position of *A. arabiensis* (now population A and B, for *296S* and *wt* respectively) and *A. coluzzii* populations (now population C, together with the *A. merus* negative control). Under this setup, we expect to see evidence of introgression between *296S A. coluzzii* and *296S A. arabiensis* in both tests (positive controls), but a positive result with any of the *wt* comparisons can indicate that *296S* haplotypes in either species is more similar to *wt* from the other (and hence, the second species is the species of origin). A detailed account of all comparisons, populations and complete statistical reports are available in Supplementary Material SM12.

We performed the same series of tests for the *A. gambiae* ∼ *A. coluzzii* introgression of the *296G* cluster in individuals without the 2La inversion (2L+^a^/2L+^a^ homozygotes; Figure 7B and Supplementary Material SM13A, B) and with the 2La inversion (Supplementary Material SM13C, D). In these tests, homozygous individuals from various *A. gambiae* and *A. coluzzii* populations were alternatively used as groups A/B (A if *296G*, B if *wt*) and C (*296G* and *wt*, separately); and *wt* outgroups were selected according to their 2La karyotype (2L+^a^/2L+^a^: *A. quadriannulatus* and *A. melas*; 2La/2La: *A. merus*). A detailed account of all comparisons, populations and complete statistical reports are available in Supplementary Material SM13.

### Sequence divergence between 2La karyotypes

To ascertain whether *296G* karyotypes from 2La chromosomes were introgressed from a 2L+^a^ background, we calculated the absolute sequence divergence (*Dxy* (Takahata and Nei 1985)) around the *Rdl* locus between all combinations of the following groups of haplotypes: (i) between *296G*-carrying haplotypes from 2L+^a^/2L+^a^ homozygotic genomes, (ii) *wt* haplotypes from 2La/2La; (iii) *296G* haplotypes from 2La/2La, (iv) *wt* haplotypes from 2La/2La (Figure 8). *Dxy* estimates were calculated along the 2L arm using the *windowed_divergence* utility in *scikit-allel* (window size=20,000 bp with 10% overlap). At each window, we also calculated the ratio between the following *Dxy* estimates: (i) *296G*-2L+^a^ ∼ *wt*-2La / *296G*-2L+^a^ ∼ *wt*-2L+^a^; and (ii) *296G*-2La ∼ *wt*-2La / *296G*-2La ∼ *wt*-2L+^a^. Thus, windows with ratios >1 are more similar to the *wt*-2L+^a^ background, and windows with ratios <1 are more similar to the *wt*-2La background.

### Alignment of *Rdl* orthologs

We retrieved *Rdl* orthologs from the following species of the Culicidae family (available in Vectorbase): *A. gambiae*, *A. arabiensis*, *A. melas*, *A. merus*, *A. christyi*, *A. epiroticus*, *A. minimus*, *A. culicifacies*, *A. funestus*, *A. stephensi*, *A. maculatus*, *A. farauti*, *A. dirus*, *A. atroparvus*, *A. sinensis*, *A. albimanus*, *A. darlingi*, *Aedes aegypti*, *Aedes albopictus*, and *Culex quinquefasciatus*. We retained (i) those orthologs that resulted in complete predicted peptides (defined as having the same start and end codons as the *A. gambiae Rdl*), and (ii) the longest isoform per gene (except for *A. gambiae*, where all three isoforms were retained). These sequences were aligned using *MAFFT* 7.310 (1,000 rounds of iterative refinement, G-INS-i algorithm) (Katoh and Standley 2013). Pairwise sequence identity between peptide sequences was calculated using the *dist.alignment* function (with a identity distance matrix, which was then converted to a pairwise identities) from the *seqinr* 3.4-5 library (Charif and Lobry 2007), in R 3.6.1 (R Core Team 2017). Pairwise *d_N_*/*d_S_* ratios were calculated from a codon-aware alignment of CDS sequences, using the *dnds* function from the *ape* 5.3 R library (Paradis et al. 2004). The codon-aware alignment of full-length CDS was obtained with PAL2NAL (Suyama et al. 2006), using the peptide alignment as a reference. Tables of pairwise identity and *d_N_*/*d_S_* values have been created with *pheatmap* 1.0.12 (Kolde 2019).

### Homology modelling and automated ligand docking

The structure of human GABA_A_ receptor bound with picrotoxin (PDB accession: 6HUG) provided the template for generating a homology model of the homopentameric *A. gambiae* RDL receptor (UniProtKB accession: Q7PII2). Sequences were aligned using *Clustal Omega* (Sievers et al. 2011), and 50 homology models were generated using *MODELLER* 9.23 (Eswar et al. 2006). A single best model was chosen based on the internal scoring values from *MODELLER* and by visually inspecting models in *Swiss-PdbViewer* (Guex et al. 1999) to eliminate candidates with structural problems. The *A296G* and *A296S* mutants were generated using *Swiss-PdbViewer* to introduce the amino acid substitutions and to energy minimise the resulting structures using 50 steps of conjugate gradient energy minimization. The pore radii of the channel models were calculated using *HOLE* 2.0 (Smart et al. 1996). The 3-dimensional structure of dieldrin was generated ab initio using *MarvinSketch* 19.22 of the ChemAxon suite (ChemAxon 2019). *AutoDockTools* 1.5.6 (Morris et al. 2009) was used to define rotatable bonds and merge non-polar hydrogens. Automated ligand docking studies with the wild-type GABA receptor model were performed using *AutoDock Vina* 1.1.2 (Trott and Olson 2009) with a grid of 20 × 20 × 20 points (1Å spacing) centred on the channel pore. Figures were produced using *PyMOL* (Schrödinger 2015).

### Availability of code and data

Python (3.7.4) and R scripts (3.6.1) to reproduce all analyses in this manuscript are available on GitHub: https://github.com/xgrau/rdl-Agam-evolution

All genome variation data has been obtained from the publicly available repositories of the *Ag1000G* project Phase 2-AR1 (The *Anopheles gambiae* 1000 Genomes Consortium 2017). Accession codes are available in Supplementary Material SM1 and download instructions can be found in the above-mentioned GitHub repository.

## Supporting information

Supplementary Material SM1. Data sources

Supplementary Material SM2. List of genetic variants in Rdl

Supplementary Material SM3. Linkage disequilibrium in Rdl

Supplementary Material SM4. Alignments of Rdl orthologs

Supplementary Material SM5. Haplotype classification and population frequency

Supplementary Material SM6. Minimum spanning networks of Rdl haplotypes

Supplementary Material SM7. Linkage disequilibrium of Rdl and Vgsc

Supplementary Material SM8. Co-segregation of Rdl and Vgsc mutations

Supplementary Material SM9. PCA of 2La karyotypes

Supplementary Material SM10. Alignments of Rdl haplotypes

Supplementary Material SM11. Phylogenies of Rdl haplotypes

Supplementary Material SM12. 296S introgression between A. coluzzii and A. arabiensis

Supplementary Material SM13. 296G introgression between A. gambiae and A. coluzzii

Supplementary Material SM14. Diversity of 296G haplotypes in 2L+a and 2La backgrounds

Supplementary Material SM15. Genetic differentiation in the 2La inversion

## Acknowledgements

We thank Arjèn Van ‘t Hof and Eric Lucas (LSTM) for fruitful discussions on the manuscript and its methods. We also thank Chris Clarkson (Wellcome Sanger Institute) for making his code publicly available.

This work was supported by the National Institute of Allergy and Infectious Diseases (R01-AI116811; the Wellcome Trust (090770/Z/09/Z; 090532/Z/09/Z; 098051); the Medical Research Council UK and the Department for International Development (MR/M006212/1) and the Medical Research Council (MR/P02520X/1). The latter grant is a UK funded award and is part of the EDCTP2 programme supported by the European Union. The content of this manuscript is solely the responsibility of the authors and does not necessarily represent the official views of the National Institute of Allergy and Infectious Diseases, or the National Institutes of Health.

## Author contributions

XGB, MD and DW designed the study. XGB carried out the analyses of sequence diversity, selection and introgression, with assistance and code contribution from ST, NJH and AM. AOR carried out the structural modelling analyses. The *Ag1000G* Consortium undertook collection, preparation, sequencing, and primary analysis of the samples. All authors read and approved the final manuscript.

## Competing interests

The authors declare no competing interests.

## Supplementary legends

**Supplementary Material SM1. Data sources.** List of genome samples from *Ag1000G* Phase 2-AR1 (table A), the Phase 1-AR3 subset (table B) (both of which contain *A. gambiae* and *A. coluzzii* specimens), and outgroup species (table C; includes *A. arabiensis*, *A. quadriannulatus*, *A. christyi*, *A. epiroticus*, *A. merus* and *A. melas*). For each sample, we include their country and population of origin, accession numbers (based on *Ag1000G* for Phase 1 and 2, and on NCBI SRA for outgroups), and the estimated 2La karyotypes.

**Supplementary Material SM2. List of genetic variants in *Rdl.* A)** List of all variants present in the *Rdl* gene (AGAP006028), including their genomic coordinates, reference and alternative alleles, coordinates of the mutation along *Rdl* CDS and peptide sequences, effect on the peptide sequence (aminoacid substitution), and frequencies in each of the populations of the cohort (Phase 2 and outgroups). **B)** Genotypes of *Rdl* non-synonymous mutations in each sample (for the six mutations reported in Figure 1), where 0=wt homozygote, 1=heterozygote, 2=alternate allele homozygote.

**Supplementary Material SM3. Linkage disequilibrium in *Rdl.*** Linkage disequilibrium between non-synonymous mutations in *Rdl*, separated by population. Only populations where non-synonymous variants are shown are displayed. For each population, we display Huff and Rogers’ *r* (left) and Lewontin’s *D′* (right).

**Supplementary Material SM4. Alignments of *Rdl* orthologs. A)** Alignment of *Rdl* orthologs from 12 species from the Culicidae family: *A. gambiae* (Anogam) *A. arabiensis* (Anoara), *A. atroparvus* (Anoatr), *A. darlingi* (Anodar), *A. dirus* (Anodir), *A. epiroticus* (Anoepi), *A. farauti* (Anofar), *A. funestus* (Anofun), *A. merus* (Anomer), *A. minimus* (Anomin), and *Aedes aegypti* (Aedaeg). Pfam-predicted protein domains, transmembrane regions and the 296 and 345 codons are shown on top of the alignment (coordinates based on the *A. gambiae* ortholog). **B-C)** Pairwise sequence identity and *d_N_/d_S_* between *Rdl* orthologs, including all *A. gambiae* isoforms (RA, RB, RC).

**Supplementary Material SM5. Haplotype classification and population frequency. A)** Clustering of haplotypes according to the minimum spanning networks (built from 626 phased variants located around codon 296; Figure 3 and Supplementary Material SM6). For each cluster, we report their population and country of origin, species, and allele present in *Rdl* codon 296 (*296G*, *296S*, *wt*) and *Vgsc* codon 995 (*995F*, *995S* and *wt*). Cluster “4” includes haplotypes with *296G* alleles, cluster “34” includes *296S* alleles; all other clusters are *wt*. **B)** Absolute frequency of *296G*, *296S* and *wt* haplotype clusters per population.

**Supplementary Material SM6. Minimum spanning networks of *Rdl* haplotypes.** Minimum spanning networks of haplotypes around *Rdl* codon 296 (626 phased variants located +/− 10,000 bp from the 2L:25429236 position), including all non-singleton haplotype clusters. Purple arrows indicate the direction of non-synonymous mutations (relative to reference assembly). **A)** Nodes are color-coded according to genotype in *Rdl* codon 296 . **B)** Nodes are color-coded according to genotype in *Vgsc* codon 995. **C)** Nodes are color-coded according to species.

**Supplementary Material SM7. Linkage disequilibrium of *Rdl* and *Vgsc*.** Linkage disequilibrium between non-synonymous mutations in *Rdl* and *Vgsc*, calculated using Huff and Rogers’ *r* (A) and Lewontin’s *D′* (B). Resistance variants in both genes are highlighted in orange (*Vgsc*) and cerise red (*Rdl*).

**Supplementary Material SM8. Co-segregation of *Rdl* and *Vgsc* mutations. A-B)** Frequency of alleles in *Vgsc* codon 995 and *Rdl* codon 296 per population, calculated per chromosome. Note: *A. gambiae* populations denoted with an asterisk (The Gambia, Guinea-Bissau and Kenya) are listed separately due to their high frequency of hybridisation and/or unclear species identification (see Methods). **C)** Geographical co-occurrence of *Rdl* and *Vgsc* mutations, at 10% and 30% frequency thresholds (chosen for illustrative purposes). Dots indicate presence. **D)** Euler diagrams and contingency table depicting the co-occurrence of *Vgsc 995F* and *995S* alleles with *Rdl 296G*, *296S* and *wt* alleles within chromosomes analysed in this study (*n* = 2356). For chromosomes carrying each of the *Rdl* haplotype groups, we include the percentage of associated genotypes at *Vgsc* codon 995. **E)** Number of chromosomes carrying *296S* or *296G* mutations (x axis) against number of *995F* mutations (y axis), per population (only values >0 included). **F)** Contingency tables of *Rdl* and *Vgsc* resistance mutations co-occurrence, per population. Only populations were resistance alleles in are segregating in both genes are included. *p* values and odds ratios [OR] correspond to Fisher’s exact tests (one-sided, testing for a greater co-occurrence of *Rdl* codon 296 and *Vgsc* 995 resistance alleles).

**Supplementary Material SM9. PCA of 2La karyotypes.** Principal component (PC) analysis of allele presence/absence from 10,000 random variants located within the 2La inversion (coordinates: 2L:20524058-42165532). Specimens from *Ag1000G* Phase 1 and *A. arabiensis* are color-coded by 2La genotype (homozygotes and heterozygotes, blue-purple), and they are used as a reference to assign 2La genotypes to Phase 2 specimens (grey). Panels A and B show PC1, PC2 and PC3; panel C shows the fraction of variance explained by each PC. The 2La karyotypes of all Phase 2 specimens are available in Supplementary Material SM1.

**Supplementary Material SM10. Alignments of *Rdl* haplotypes. A)** 5′ start of the gene (2L:25363652, 696 variants). **B)** 3′ end of the gene (2L:25434556, 428 variants). **C)** Unadmixed upstream region within the 2La inversion (1 Mb upstream of *Rdl*; 2903 variants). **D)** Unadmixed downstream region within the 2La inversion (1 Mb downstream of *Rdl*, 2594 variants). The name of each sequence name indicates the specimen (codes from Supplementary Material SM1; e.g. AA0040-C), haplotype (a or b), population of origin (e.g. GHcol), genotype at codon 296 (gt0=wt, gt1=296G, gt2=296S), and 2La background (kt0=2L+^a^/2L+^a^, kt1=2La/2L+^a^, kt2=2La/2La).

**Supplementary Material SM11. Phylogenies of *Rdl* haplotypes.** Phylogenetic trees from alignments around the *Rdl* locus (Supplementary Material SM10), in Newick format and including ultrafast bootstrap (UFBS) statistical supports. The name of each sequence (e.g. “AA0040-Ca_GHcol_gt0_kt0”) indicates the specimen (codes from Supplementary Material SM1; “AA0040-C”), chromosome (“a” or “b”), population of origin (“GHcol”), allele at codon 296 (gt0=wt, gt1=296G, gt2=296S), and 2La karyotype (kt0=2L+^a^/2L+^a^, kt1=2La/2L+^a^, kt2=2La/2La).

**Supplementary Material SM12. *296S* introgression between *A. coluzzii* and *A. arabiensis*. A)** Profile of Patterson’s *D* in 2La/2La backgrounds, using *A. coluzzii* specimens as populations A and B (*296S* and *wt*, respectively); *A. arabiensis* as population C (*296S* as positive controls, *wt* as test), *A. merus* as a negative control for population C (*wt*); and either *A. christyi* or *A. epiroticus* as outgroups (*wt*). **B)** Profile of Patterson’s *D* in 2La/2La backgrounds, using *A. arabiensis* specimens as populations A and B (*296G* and *wt*, respectively); *A. coluzzii* as population C (*296S* as positive controls, *wt* as test), *A. merus* as a negative control for population C (*wt*); and either *A. christyi* or *A. epiroticus* as outgroups (*wt*).

In all panels, the hypothesis under test can be summarised as follows: if *296S* homozygotes from species *i* show evidence of introgression with *wt* homozygotes from species *j* but not with *wt* from *i*, it means that *296S* originated in species *j*. Left plots depict the entire 2L chromosomal arm (orange lines demarcate 2La inversion), and rightmost plots focus on the *Rdl* locus (*Rdl* gene coordinates highlighted in red). *D* was calculated in sliding blocks of 10,000 phased variants (with 20% overlap). For each comparison, we report the mean value of *D* in the *Rdl* locus and use a block-jackknife procedure (block length = 100 variants) to estimate its standard error, a Z-score (standardized *D*) and *p-*value (that reflects deviation from the null expectation of *D* = 0).

**Supplementary Material SM13. *296G* introgression between *A. gambiae* and *A. coluzzii*. A)** Profile of Patterson’s *D* in 2L+^a^/2L+^a^ backgrounds, using *A. coluzzii* specimens as populations A and B (*296G* and *wt*, respectively); *A. gambiae* as population C (*296G* as positive controls, *wt* as test); and either *A. quadriannulatus* or *A. melas* as outgroups (*wt*). **B)** Profile of Patterson’s *D* in 2L+^a^ backgrounds, using *A. gambiae* specimens as populations A and B (*296G* and *wt*, respectively); *A. coluzzii* as population C (*296G* as positive control, *wt* as test); and either *A. quadriannulatus* or *A. melas* as outgroups (*wt*). **C)** Profile of Patterson’s *D* in 2La/2La backgrounds, using *A. coluzzii* specimens as populations A and B (*296G* and *wt*, respectively); *A. gambiae* as population C (*296G* as positive controls, *wt* as test); and *A. merus* as outgroup (*wt*). **D)** Profile of Patterson’s *D* in 2La/2La backgrounds, using *A. gambiae* specimens as populations A and B (*296G* and *wt*, respectively); *A. coluzzii* as population C (*296G* as positive controls, *wt* as test); and *A. merus* as outgroup (*wt*).

In all panels, the hypothesis under test can be summarised as follows: if *296G* homozygotes from species *i* show evidence of introgression with *wt* homozygotes from species *j* but not with *wt* from *i*, it means that *296G* originated in species *j*. Left plots depict the entire 2L chromosomal arm (orange lines demarcate 2La inversion), and rightmost plots focus on the *Rdl* locus (*Rdl* gene coordinates highlighted in red). *D* was calculated in sliding blocks of 10,000 phased variants (with 20% overlap). For each comparison, we report the mean value of *D* in the *Rdl* locus and use a block-jackknife procedure (block length = 100 variants) to estimate its standard error, a Z-score (standardized *D*) and *p-*value (that reflects deviation from the null expectation of *D* = 0).

**Supplementary Material SM14. Diversity of *296G* haplotypes in 2L+^a^ and 2La backgrounds. A)** Profile of *EHH* decay for each group of *296G* haplotypes (*296G* in 2L+^a/^2L+^a^, 2La/2L+^a^ and 2La/2La backgrounds), built from 16,623 phased variants located +/− 150,000 bp from codon 296 (2L:25429236 position). **B)** Profile of haplotypic diversity along chromosomal arm 2L (sliding blocks of 500 variants with 20% overlap). **C)** Absolute sequence divergence (*Dxy*) between *296G* alleles of 2L+^a^ background and *wt* resistance haplotypes of 2L+^a^ and 2La backgrounds. **D)** Absolute sequence divergence (*Dxy*) between *296G* alleles of 2La background and *wt* resistance haplotypes of 2L+^a^ and 2La backgrounds. All values are calculated in windows of 20,000 kbp with 10% overlap.

**Supplementary Material SM15. Genetic differentiation in the 2La inversion.** Differentiation (Hudson’s *F_ST_*) along the 2L chromosomal arm between *A. gambiae* and *A. coluzzii* species, separated by their 2La karyotype (2La/2La or 2L+^a^/2L+^a^). Panel A shows comparisons with *A. gambiae* with 2L+^a^/2L+^a^ karyotypes, and panel B for *A. gambiae* with 2La/2La karyotypes. *F_ST_* estimates have been calculated in adjacent blocks of 5,000 phased variants with 20% overlap. Sub-panels at the right focus on the *Rdl* genomic locus. Note that interkaryotype comparisons have higher *F_ST_* in the 2La region than inter-species comparisons.

## Notes

#### Summary of Updates

Updated title, abstract and introduction. No changes in the results or discussion sections.

https://github.com/xgrau/rdl-Agam-evolution

