## Supplementary figures and images for "Evolution of the insecticide target *Rdl* in African *Anopheles* is driven by interspecific and interkaryotypic introgression"

### Supplementary Material SM3. Linkage disequilibrium in Rdl

#### Supplementary Material 3

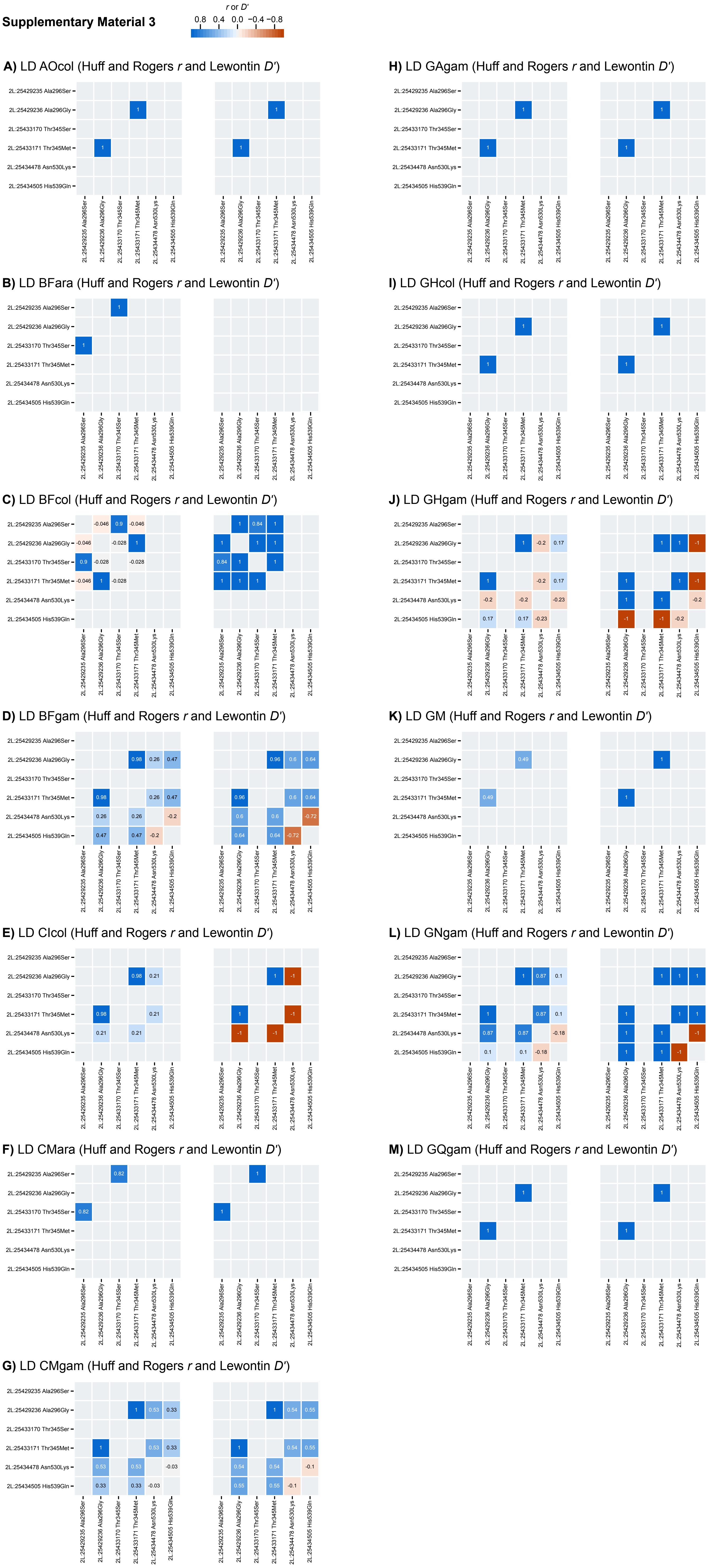

### Supplementary Material SM9. PCA of 2La karyotypes

**Supplementary Material 9**

**A)** PCA: PC1 ~ PC2

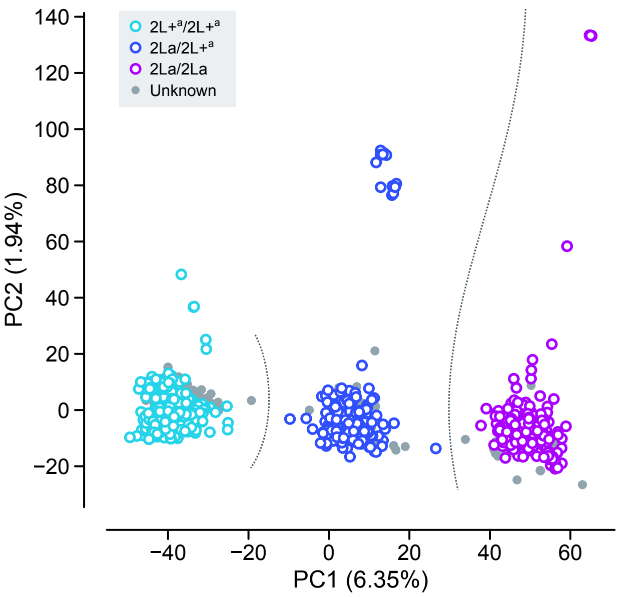


**B)** PCA: PC1 ~ PC3

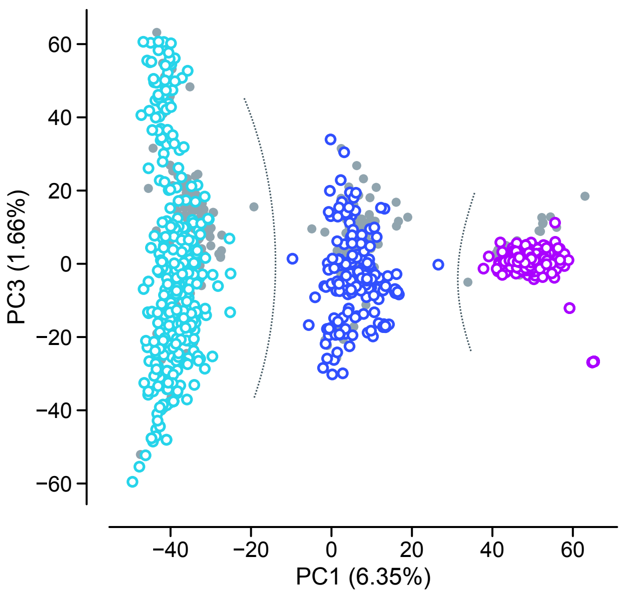


**C)** Variance explained per component

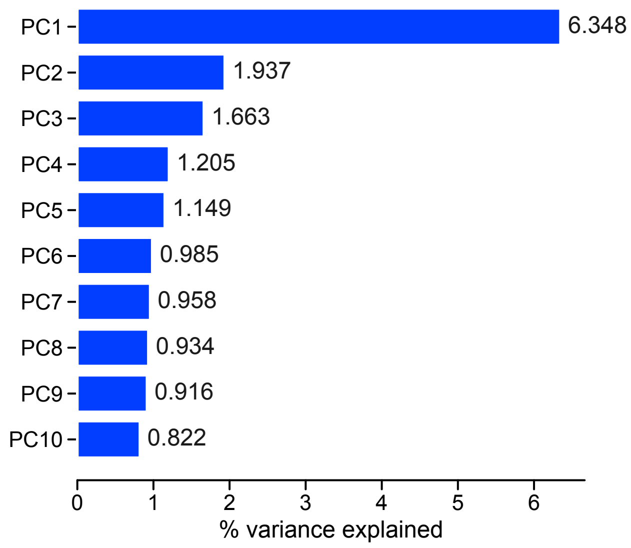

### Supplementary Material SM15. Genetic differentiation in the 2La inversion

**Supplementary Material 15**

**A)** *A. gambiae* $2L+^{a}/2L+^{a}$ ~ other

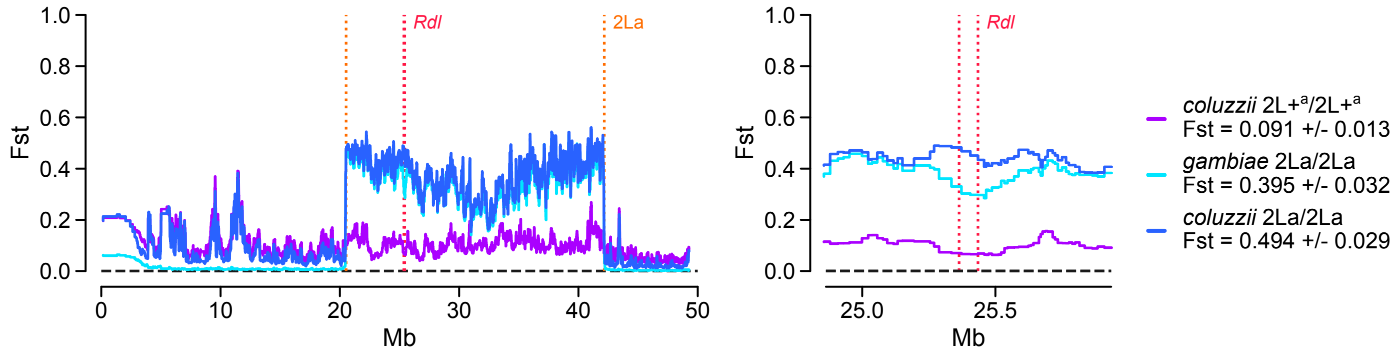


**B)** *A. gambiae* 2La/2La ~ other

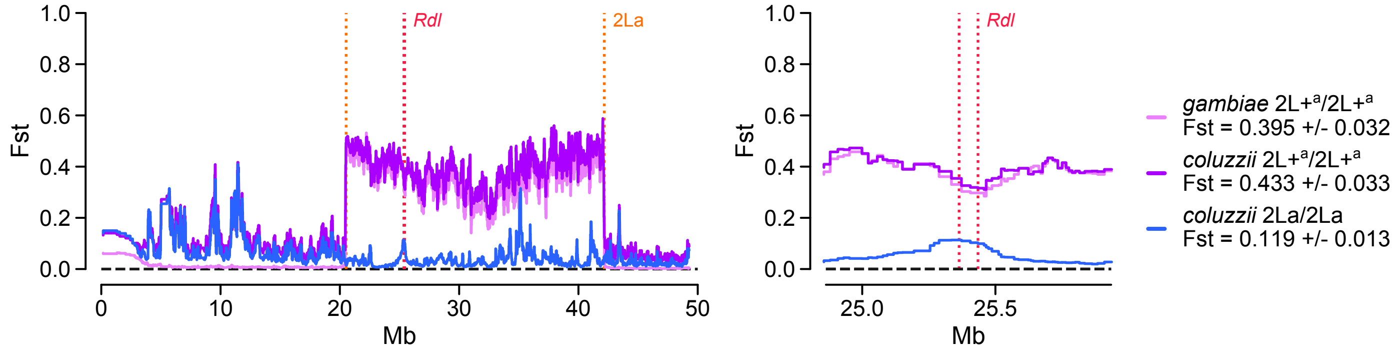
