## Supplementary Material SM4. Alignments of Rdl orthologs for "Evolution of the insecticide target *Rdl* in African *Anopheles* is driven by interspecific and interkaryotypic introgression"

**Supplementary Material 4**

**A)** Alignment of *Rdl* orthologs

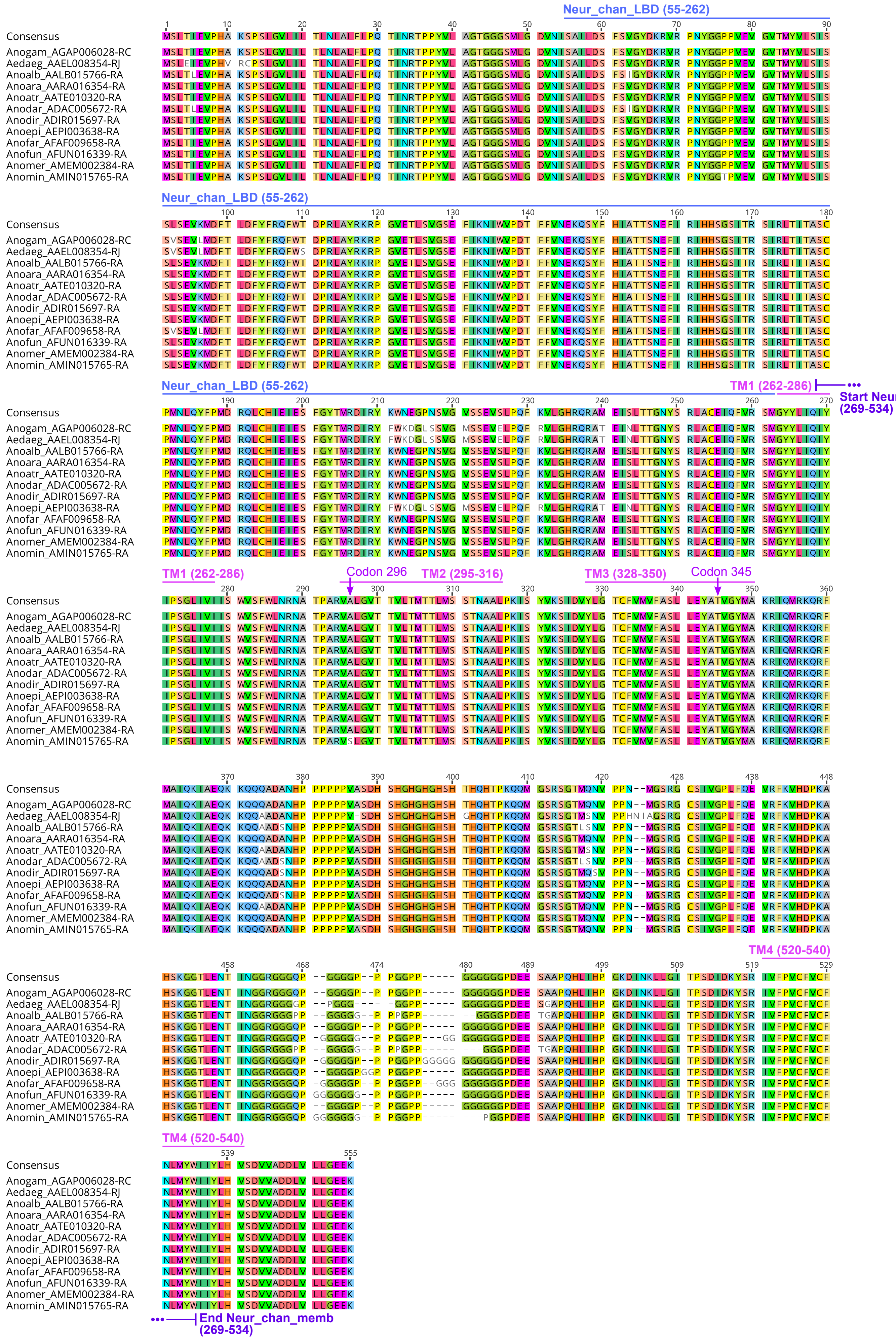


**B)** Pairwise identity (peptide alignment)

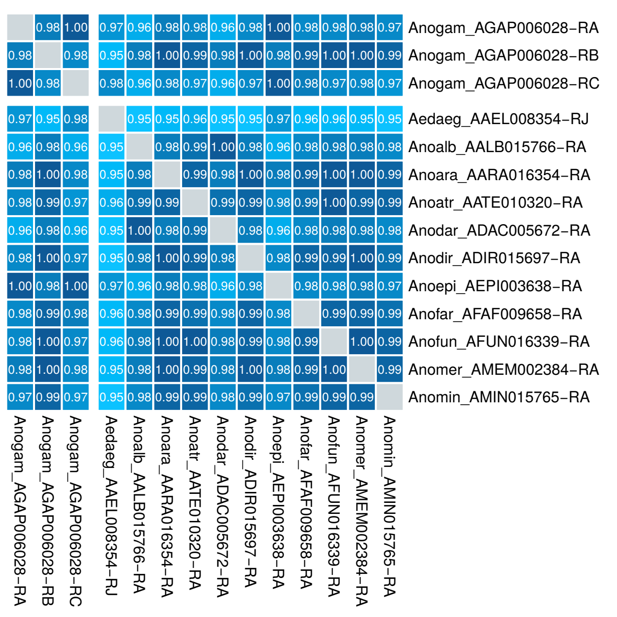


**C)** Pairwise $d_N/d_S$ (codon alignment)

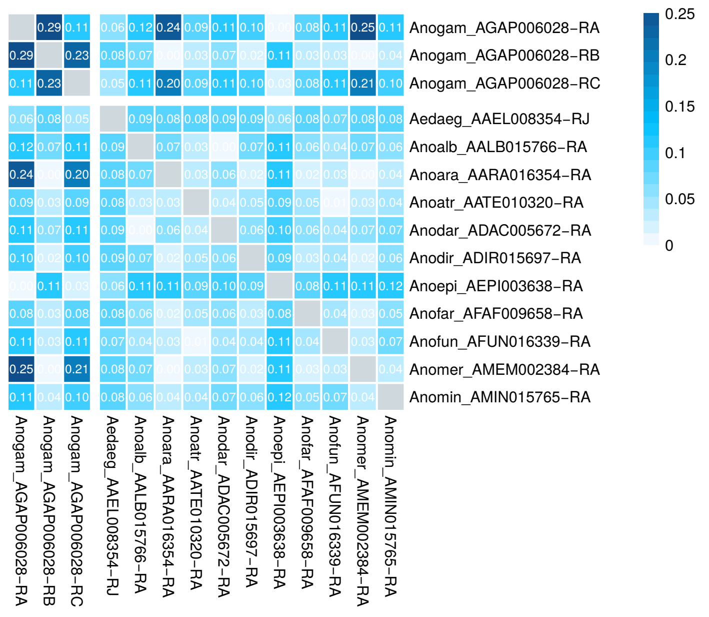
