## Supplementary Material SM6. Minimum spanning networks of Rdl haplotypes for "Evolution of the insecticide target *Rdl* in African *Anopheles* is driven by interspecific and interkaryotypic introgression"

**Supplementary Material 6**

**A)** Minimum spanning network of haplotypes in *Rdl*, colored by *Rdl* 296th codon genotype

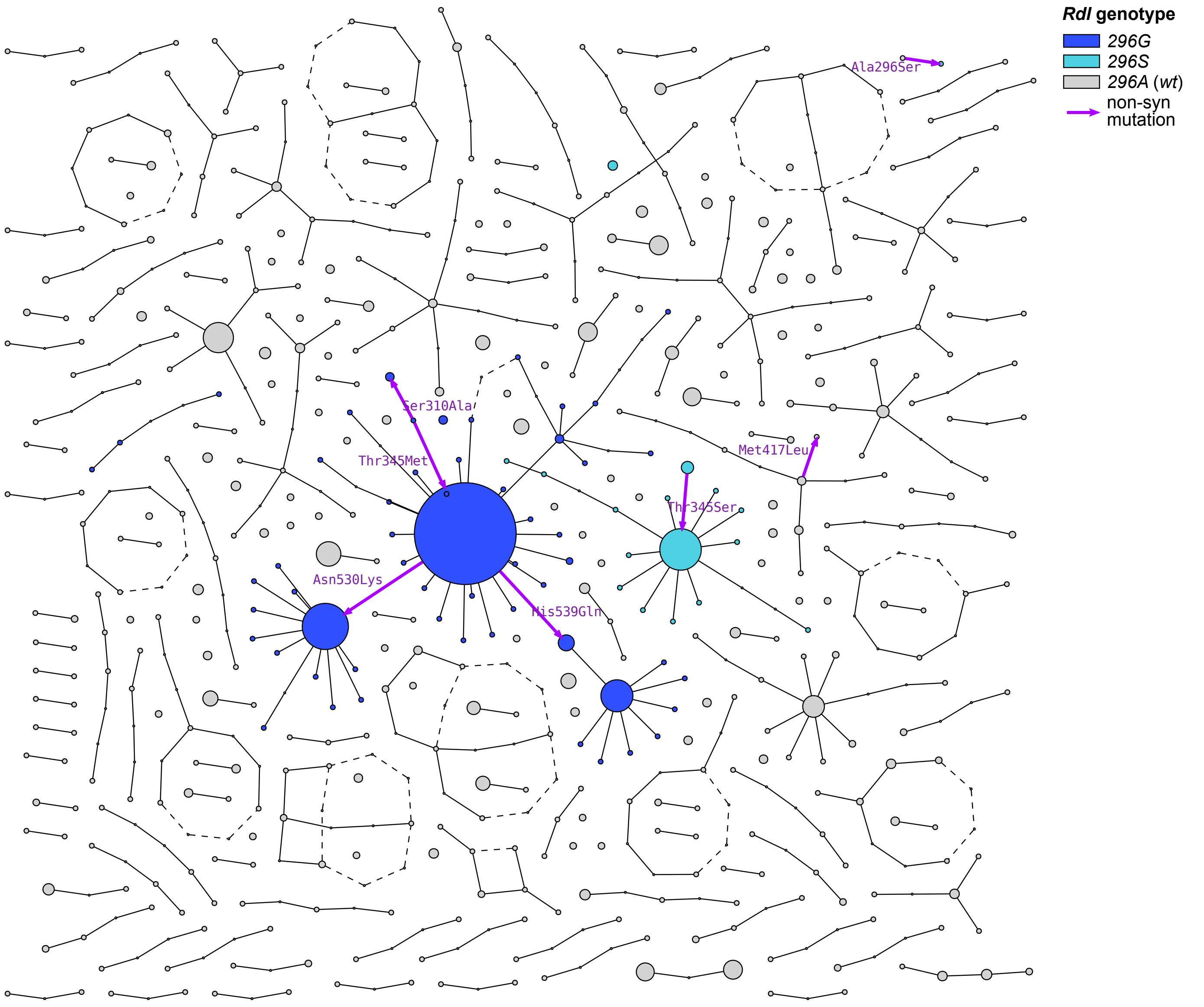


**B)** Minimum spanning network of haplotypes in *Rdl*, colored by *Vgsc* 995th codon genotype

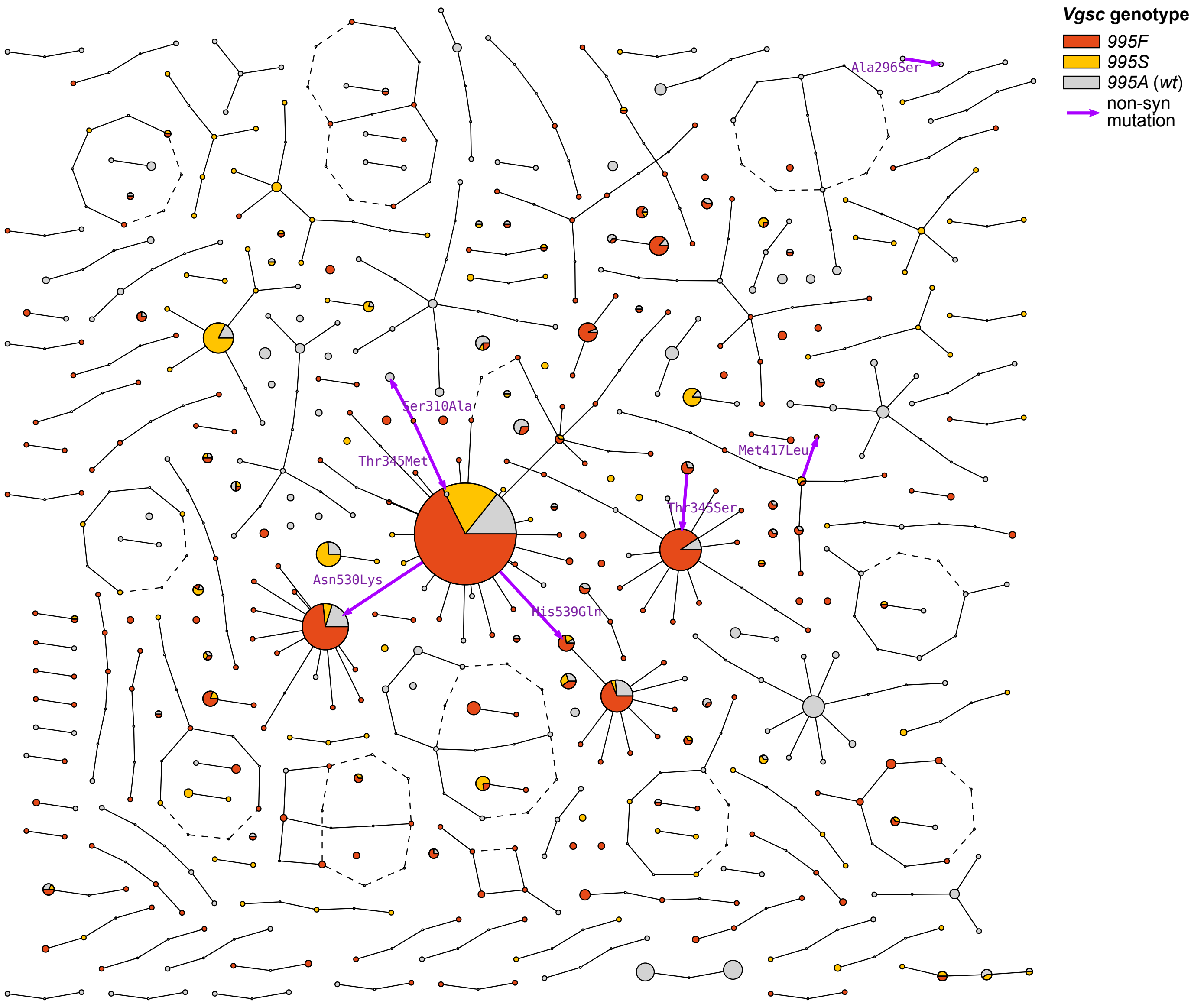


**C)** Minimum spanning network of haplotypes in *Rdl*, colored by species

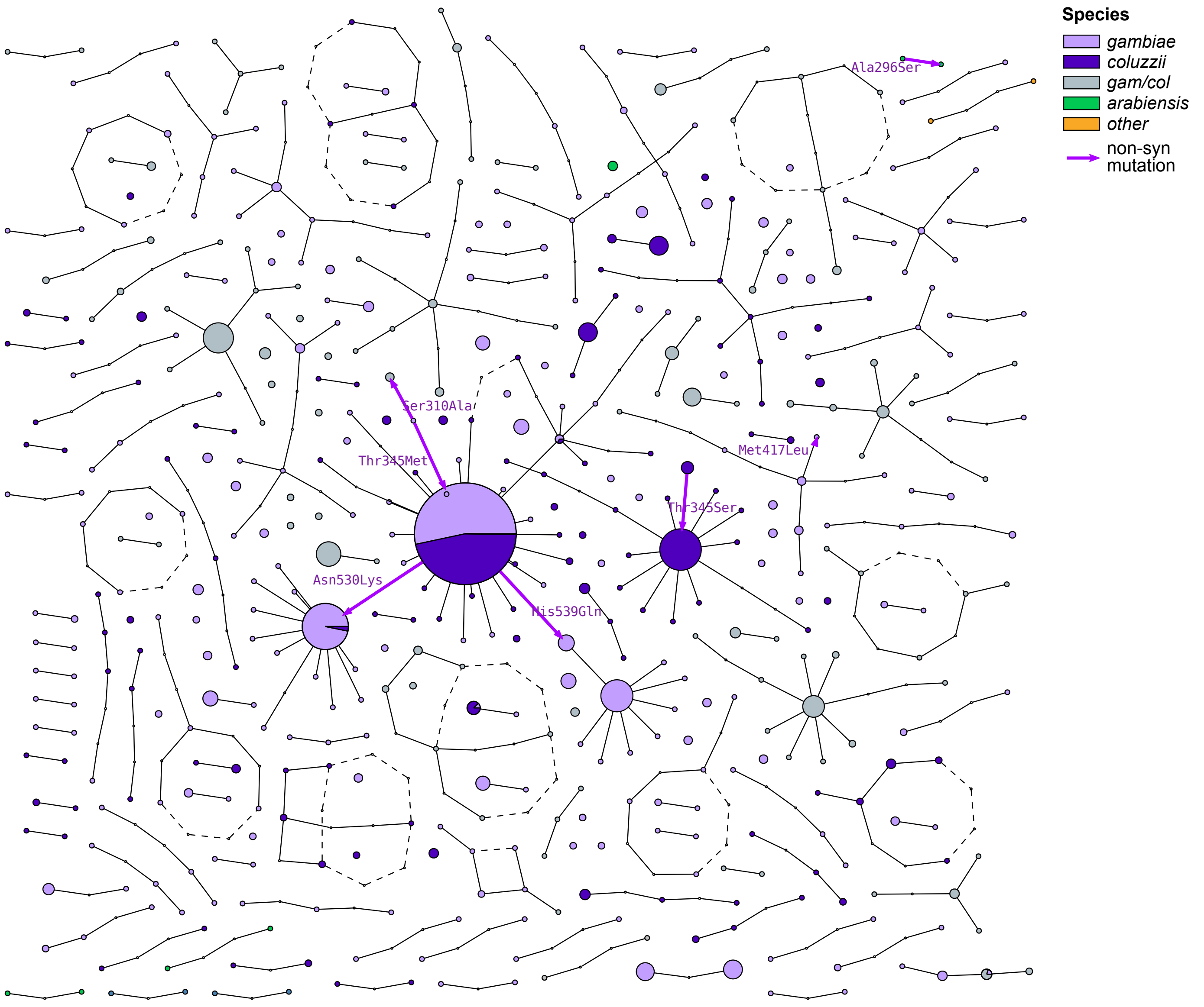
