## Supplementary Material SM7. Linkage disequilibrium of Rdl and Vgsc for "Evolution of the insecticide target *Rdl* in African *Anopheles* is driven by interspecific and interkaryotypic introgression"

**Supplementary Material 7**

**A)** Linkage disequilibrium, Huff and Rogers *r*

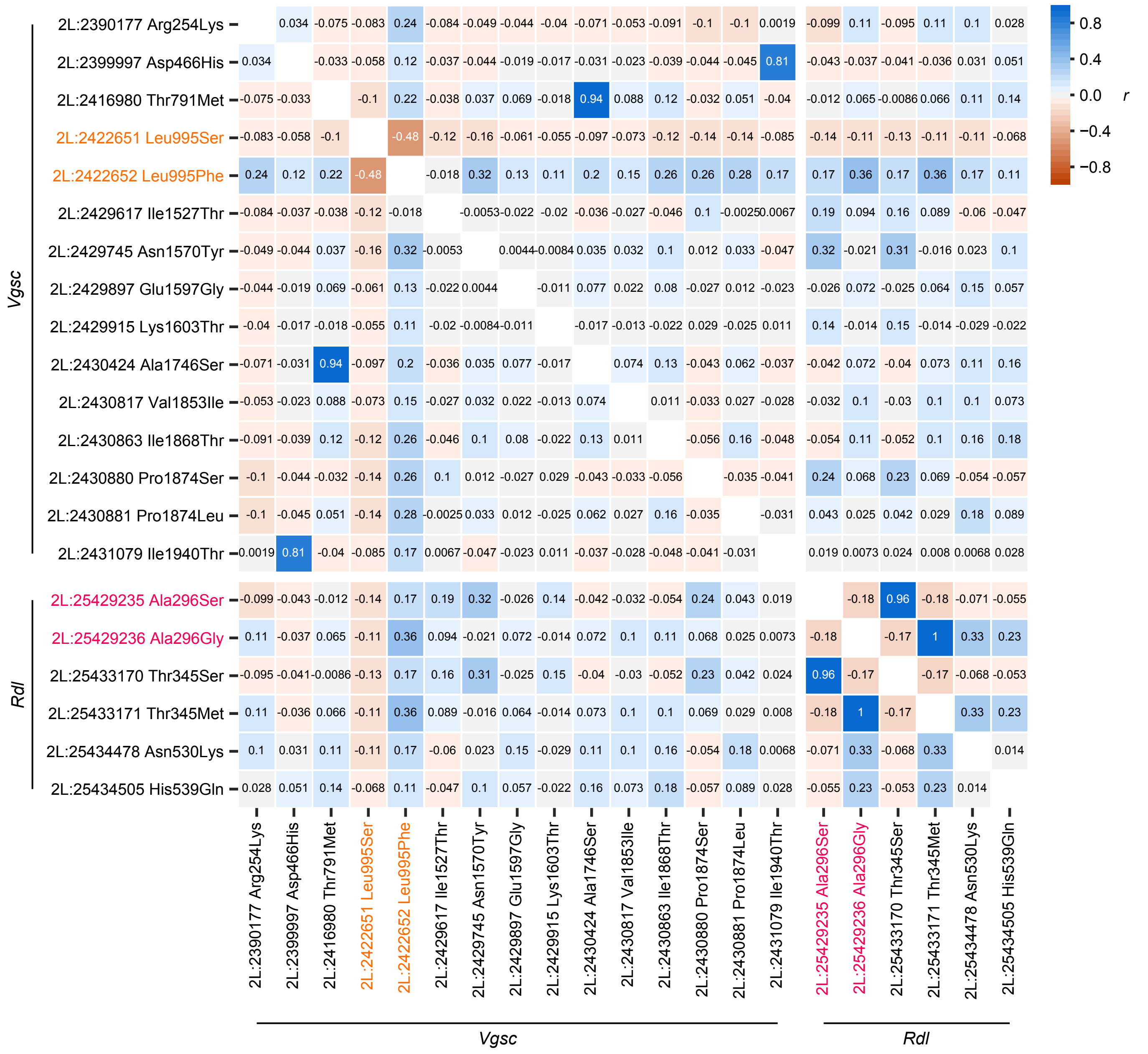


**B)** Linkage disequilibrium, Lewontin *D′*

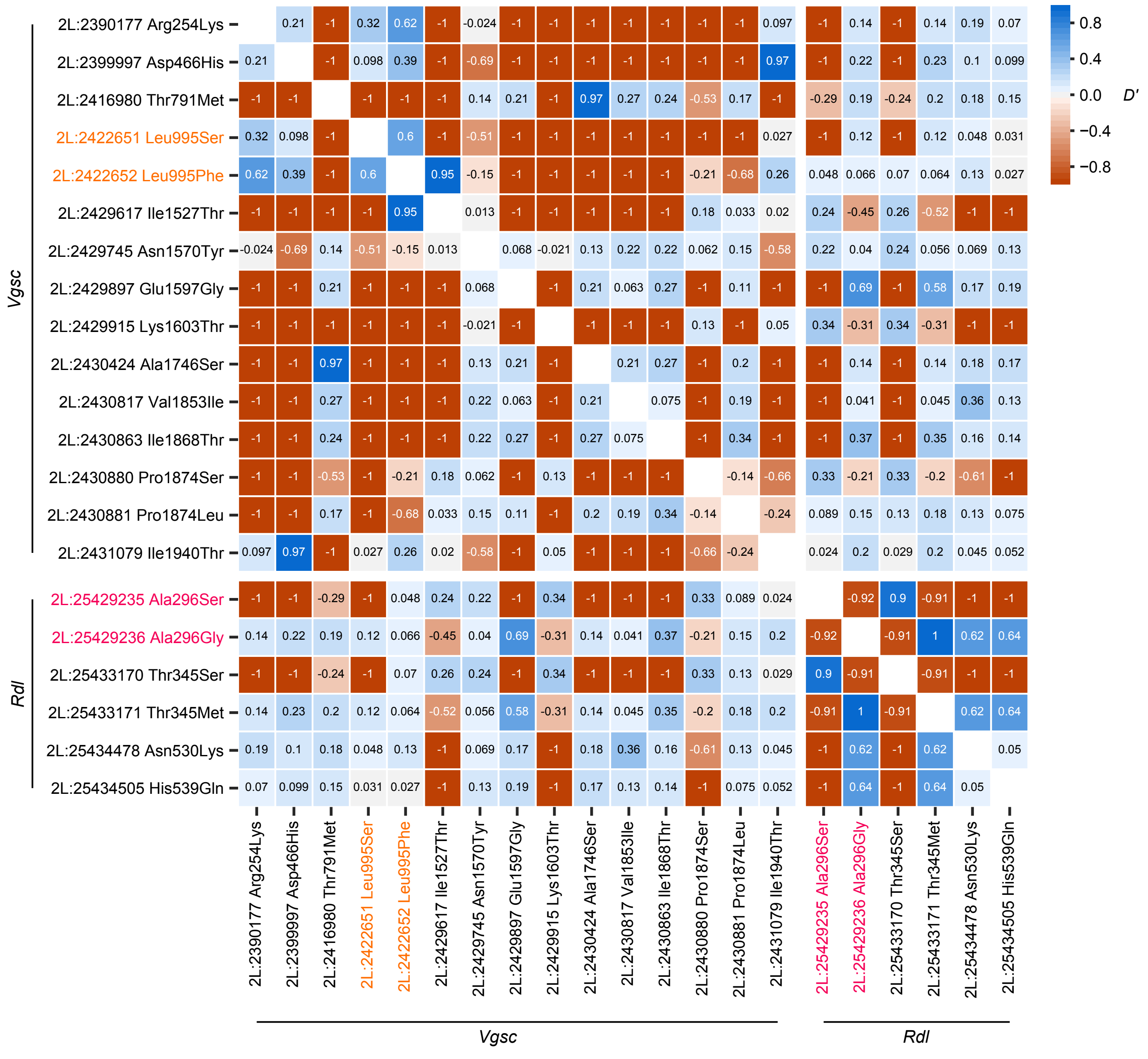
