## Supplementary Material SM8. Co-segregation of Rdl and Vgsc mutations for "Evolution of the insecticide target *Rdl* in African *Anopheles* is driven by interspecific and interkaryotypic introgression"

**Supplementary Material 8**

**A)** Frequency *Vgsc* codon 995 mutations · **B)** Frequency *Rdl* codon 296 mutations · **C)** Geographical co-occurrence

(frequency scale: 0 – 1)

| Population | A) 995F | A) 995S | A) wt | B) 296G | B) 296S | B) wt | C) 995F >30% | C) 296G/S >10% |
|---|---|---|---|---|---|---|---|---|
| Angola *A. coluzzii* | 0.84 | | 0.16 | 0.43 | | 0.57 | ● | ● |
| Burkina Faso *A. coluzzii* | 0.85 | | 0.15 | 0.01 | 0.63 | 0.37 | ● | ● |
| Côte d'Ivoire *A. coluzzii* | 0.92 | | 0.08 | 0.65 | | 0.35 | ● | ● |
| Ghana *A. coluzzii* | 0.82 | | 0.18 | 0.60 | | 0.40 | ● | ● |
| Guinea *A. coluzzii* | 0.88 | | 0.12 | | | 1.00 | ● | |
| Burkina Faso *A. gambiae* | 1.00 | | | 0.39 | | 0.61 | ● | ● |
| Cameroon *A. gambiae* | 0.53 | 0.16 | 0.32 | 0.30 | | 0.70 | ● | ● |
| Mayotte *A. gambiae* | | | 1.00 | | | 1.00 | | |
| Gabon *A. gambiae* | 0.33 | 0.67 | | 0.78 | | 0.22 | ● | ● |
| Ghana *A. gambiae* | 1.00 | | | 0.92 | | 0.08 | ● | ● |
| Guinea *A. gambiae* | 1.00 | | | 0.31 | | 0.69 | ● | ● |
| Equatorial Guinea *A. gambiae* | | | 1.00 | 0.72 | | 0.28 | | |
| Uganda *A. gambiae* | | 1.00 | | | | 1.00 | | |
| The Gambia *A. gambiae* (*) | | | 1.00 | 0.03 | | 0.97 | | |
| Guinea-Bissau *A. gambiae* (*) | | | 1.00 | | | 1.00 | | |
| Kenya *A. gambiae* (*) | | 0.76 | 0.24 | | | 1.00 | | |
| Burkina Faso *A. arabiensis* | | | 1.00 | | 0.25 | 0.75 | | |
| Cameroon *A. arabiensis* | | | 1.00 | | 0.50 | 0.50 | | |
| Tanzania *A. arabiensis* | | | 1.00 | | 0.12 | 0.88 | | |
| Zambia *A. quadriannulatus* | | | 1.00 | | 0.10 | 0.90 | | |
| South Africa *A. merus* | | | 1.00 | | | 1.00 | | |
| Kenya *A. merus* | | | 1.00 | | | 1.00 | | |
| Cameroon *A. melas* | | | 1.00 | | | 1.00 | | |

**D)** Overlap of *Rdl* codon 296 and *Vgsc* codon 995 alleles, per chromosome

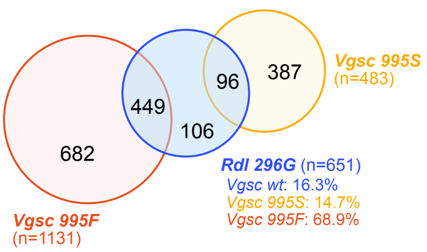


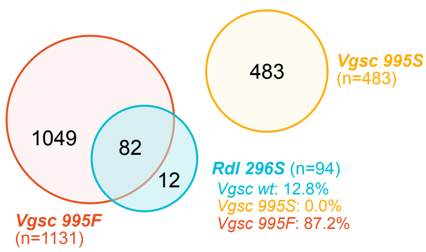


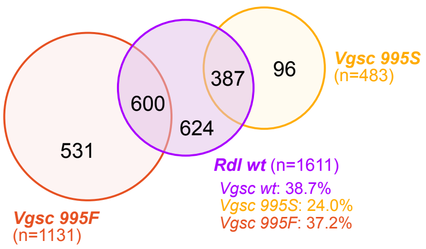


Contingency table: *Rdl* and *Vgsc* alleles

| | | *Rdl* allele 296G | 296S | wt |
|---|---|---|---|---|
| *Vgsc* allele | 995F | 449 | 82 | 600 |
| | 995S | 96 | 0 | 387 |
| | wt | 106 | 12 | 624 |
| | Total | 651 | 94 | 1611 |

**E)** Geographical co-occurrence

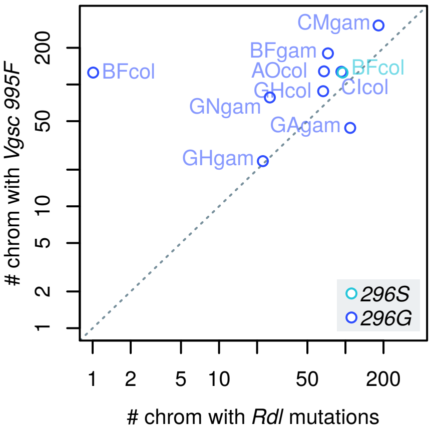


**F)** Overlap of *Rdl* codon 296 and *Vgsc* codon 995 alleles, per chromosome & populations

AOcol 995F~296G?
p = 0.0751 OR = 2.16

| | 296G | wt |
|---|---|---|
| 995F | 60 | 71 |
| wt | 7 | 18 |

BFcol 995F~296G?
p = 0.821 OR = Inf

| | 296G | wt |
|---|---|---|
| 995F | 1 | 45 |
| wt | 0 | 10 |

CIcol 995F~296G?
p = 0.965 OR = 0.344

| | 296G | wt |
|---|---|---|
| 995F | 82 | 48 |
| wt | 10 | 2 |

GHcol 995F~296G?
p = 0.898 OR = 0.589

| | 296G | wt |
|---|---|---|
| 995F | 52 | 38 |
| wt | 14 | 6 |

CMgam 995F~296G?
p = 0.444 OR = 1.05

| | 296G | wt |
|---|---|---|
| 995F | 99 | 213 |
| wt | 58 | 131 |

BFcol 995F~296S?
p = 0.252 OR = 1.51

| | 296S | wt |
|---|---|---|
| 995F | 82 | 45 |
| wt | 12 | 10 |

CMgam 995S~296G?
p = 0.838 OR = 0.786

| | 296G | wt |
|---|---|---|
| 995S | 24 | 69 |
| wt | 58 | 131 |
