## Supplementary Material SM12. 296S introgression between A. coluzzii and A. arabiensis for "Evolution of the insecticide target *Rdl* in African *Anopheles* is driven by interspecific and interkaryotypic introgression"

**Supplementary Material 12**

**A)** *A. arabiensis* as donor (C), 2La homozygotes

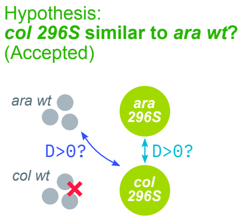


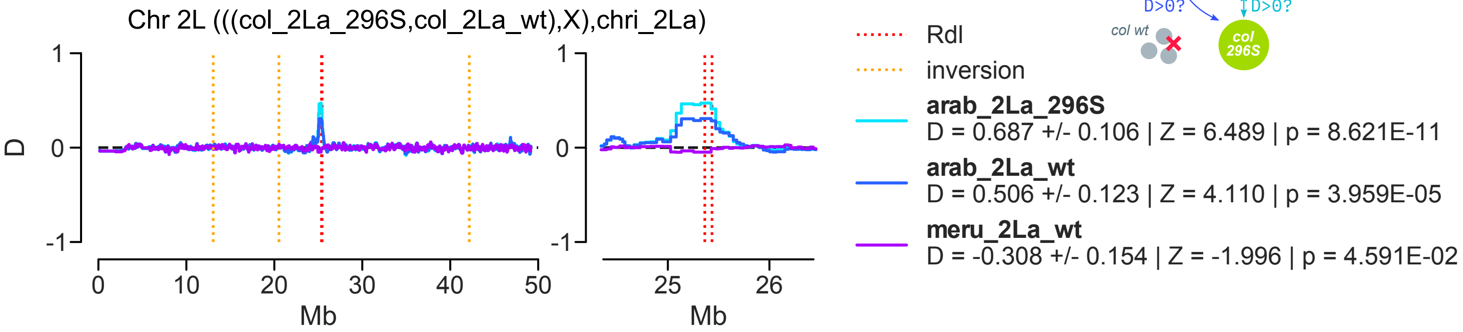


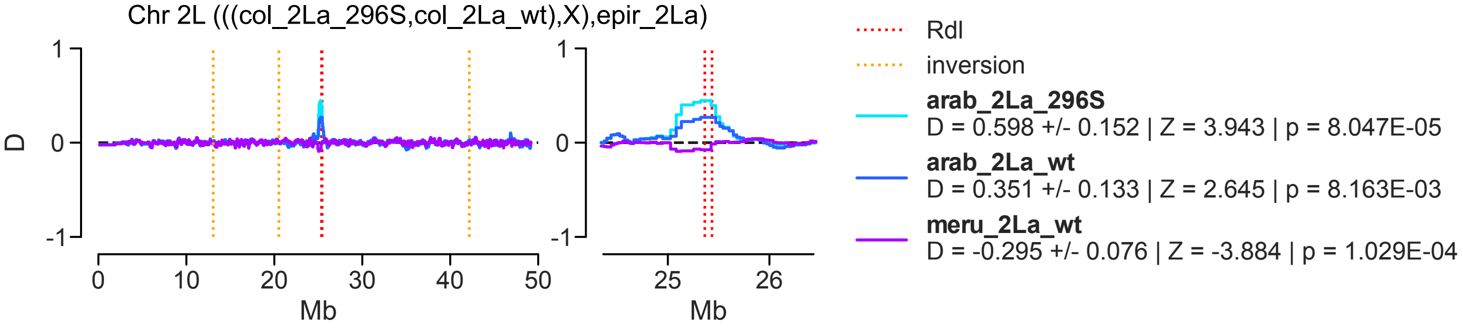


**B)** *A. coluzzii* as donor (C), 2La homozygotes

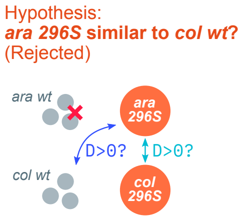


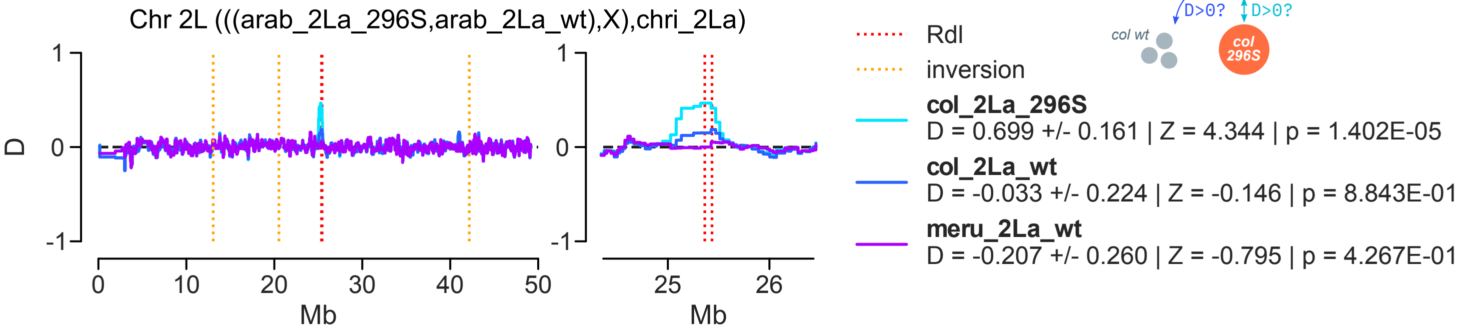


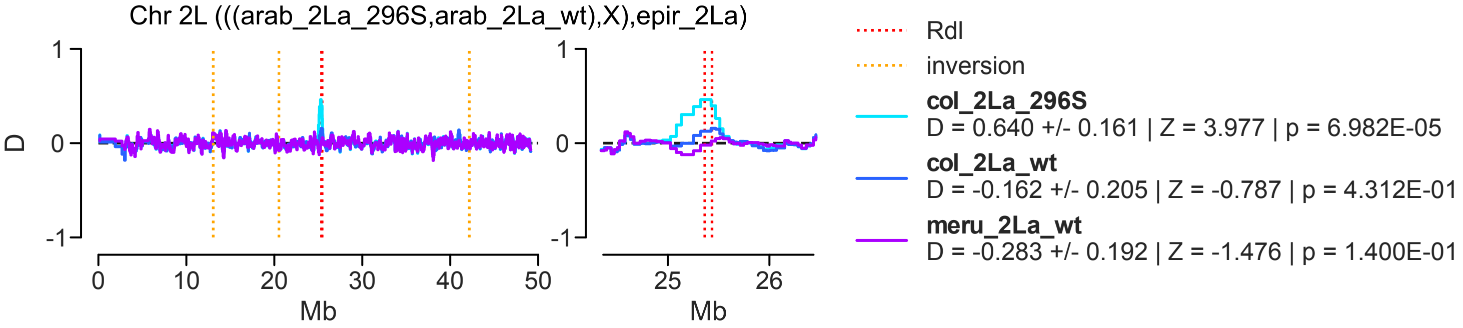
