## Supplementary Material SM13. 296G introgression between A. gambiae and A. coluzzii for "Evolution of the insecticide target *Rdl* in African *Anopheles* is driven by interspecific and interkaryotypic introgression"

Supplementary Material 13

**A)** *A. gambiae* as donor (C), 2L+ᵃ homozygotes

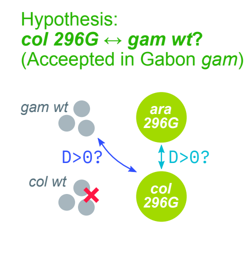


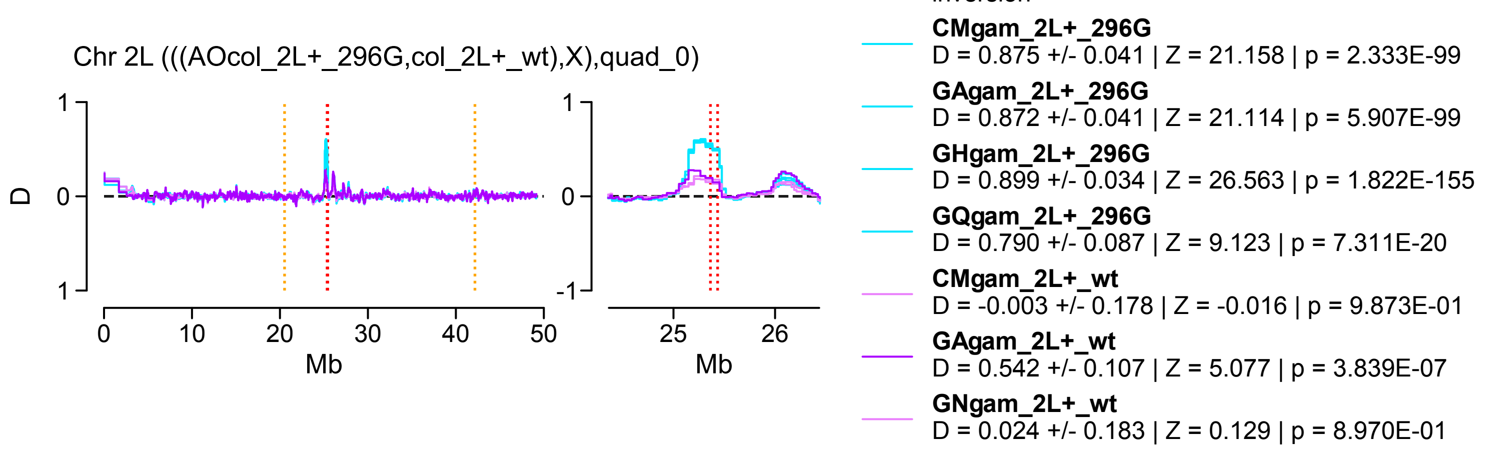


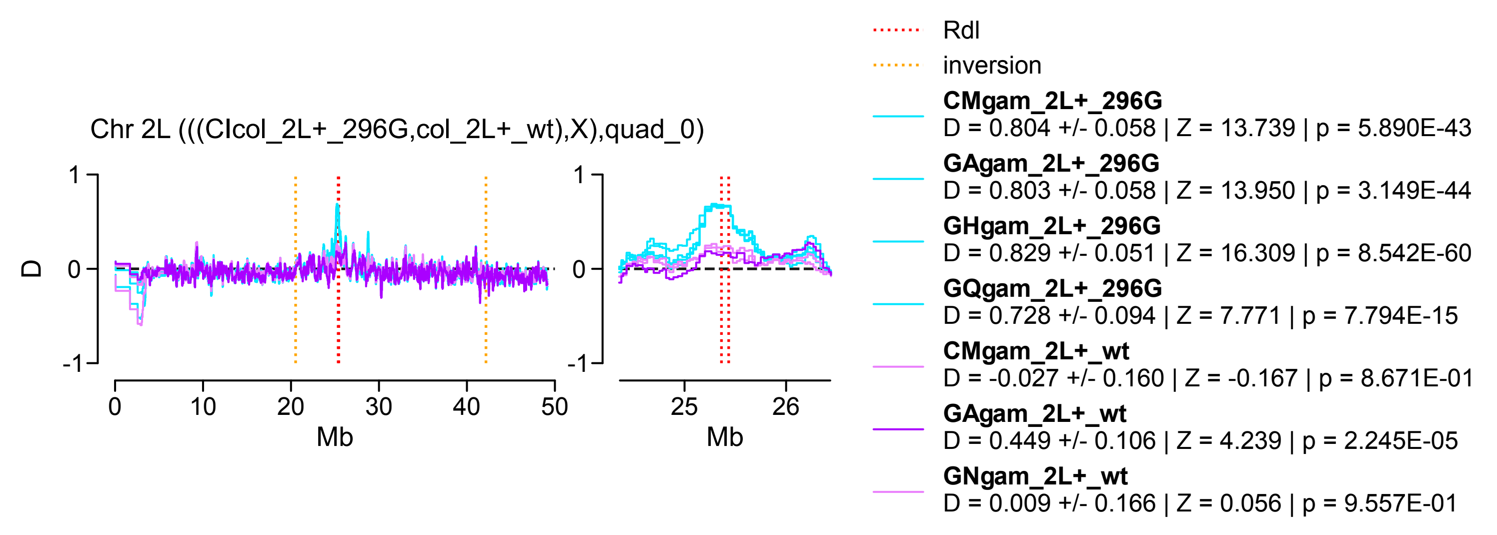


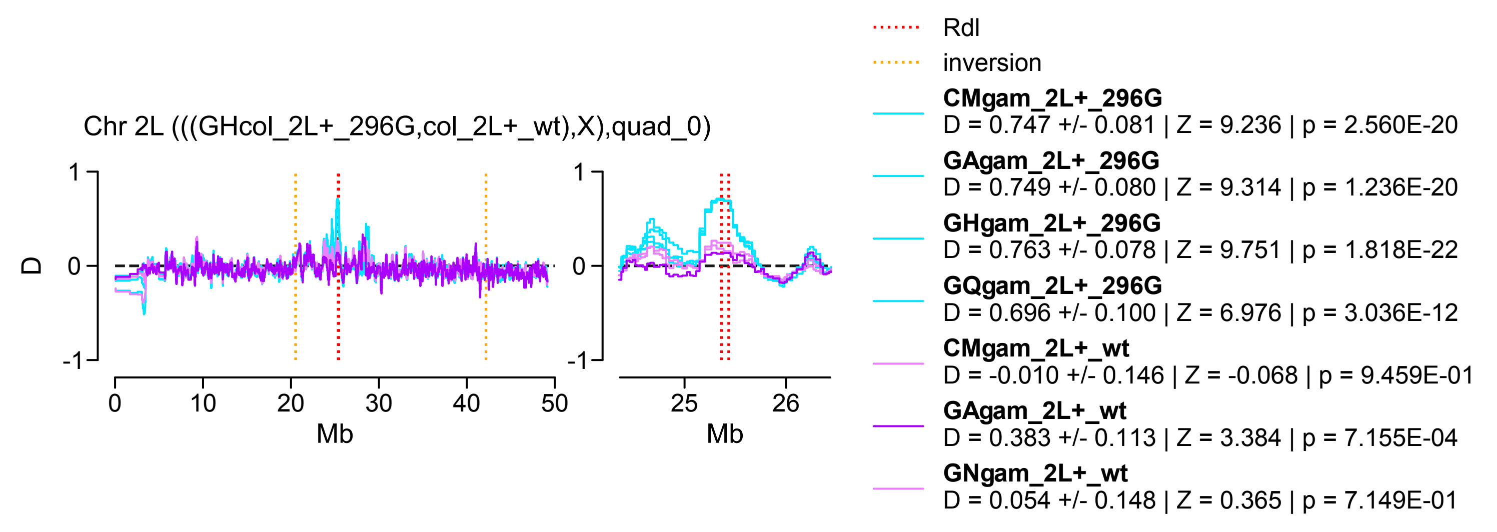


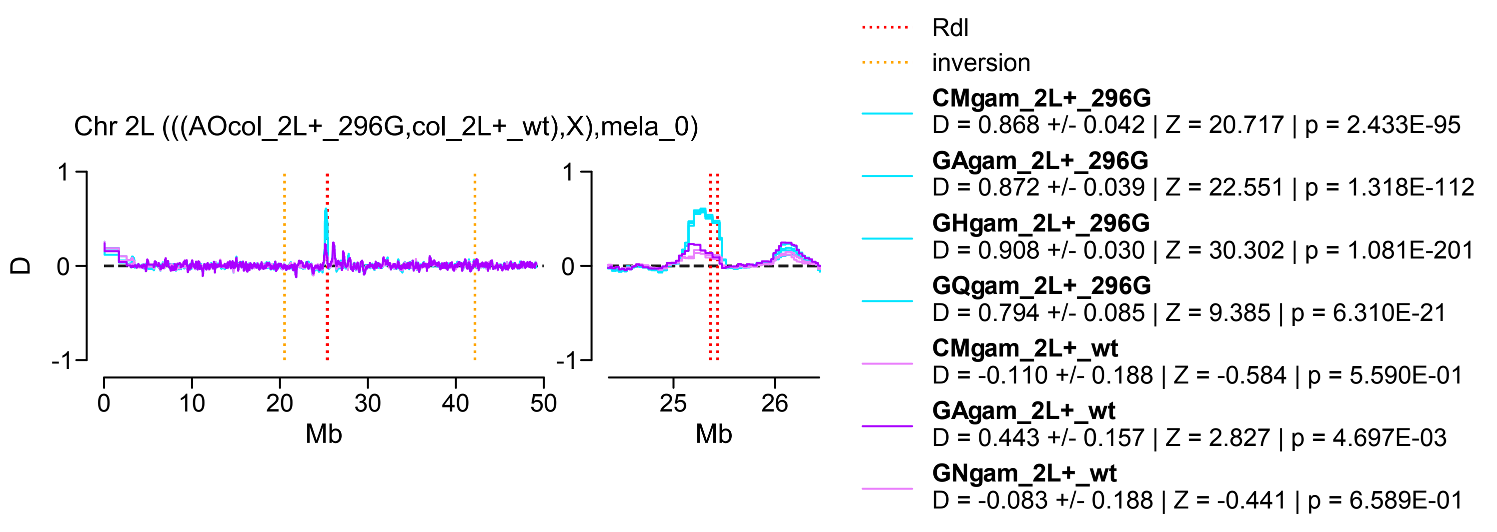


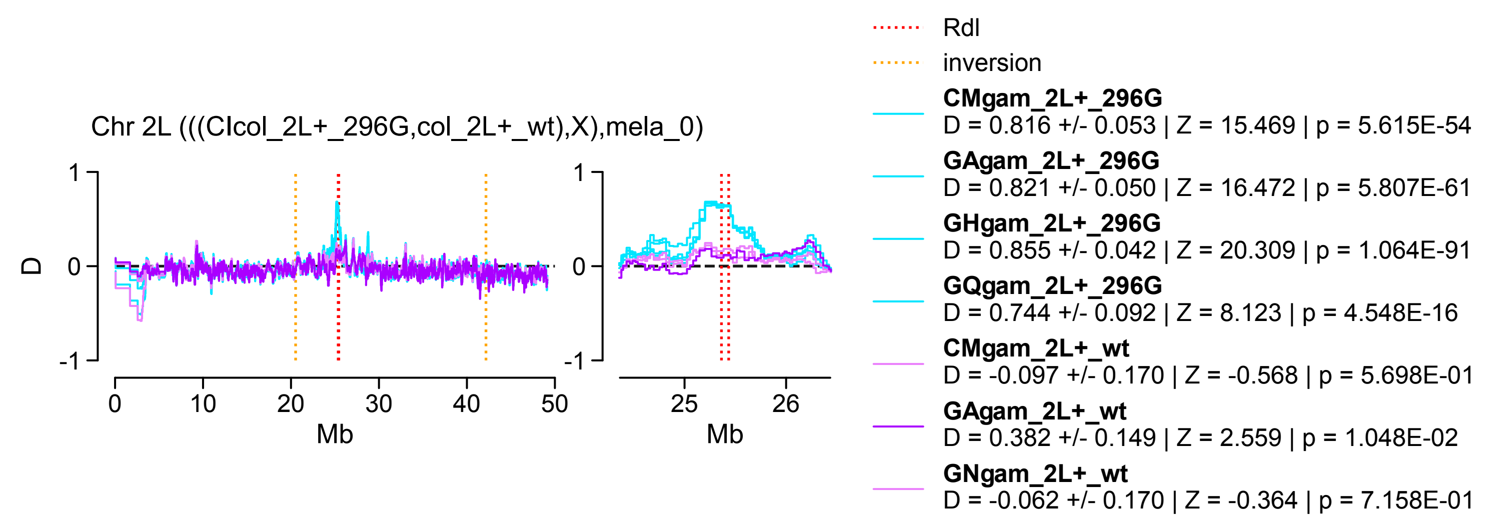


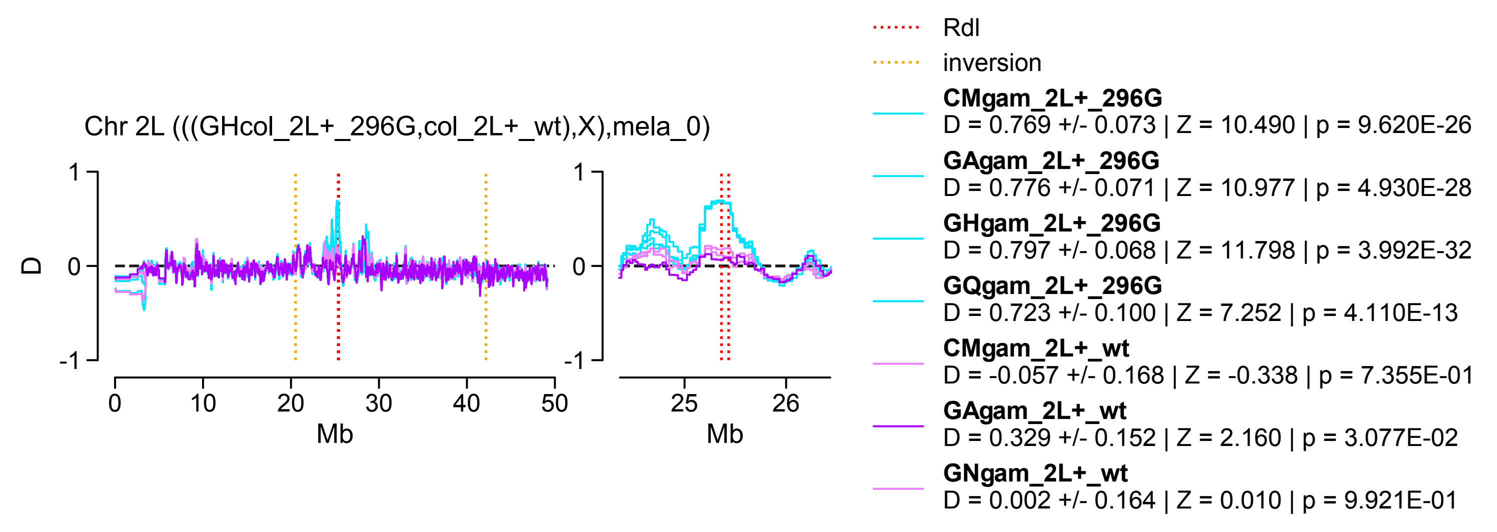


**B)** *A. coluzzii* as donor (C), 2L+ᵃ homozygotes

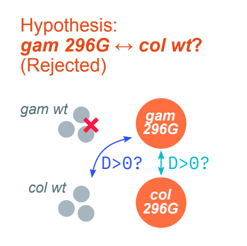


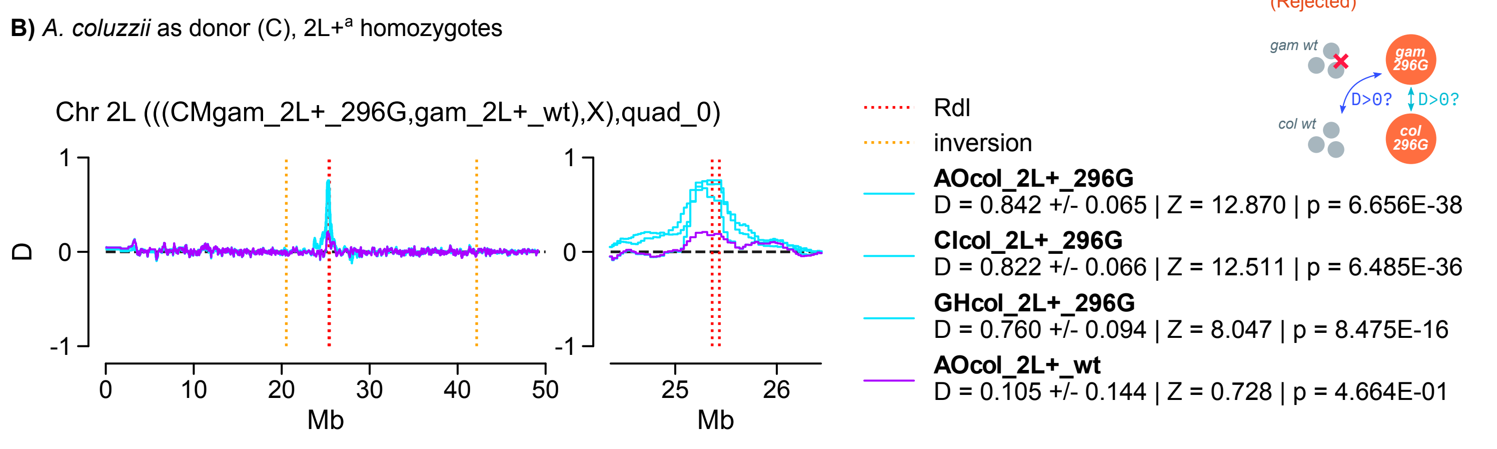


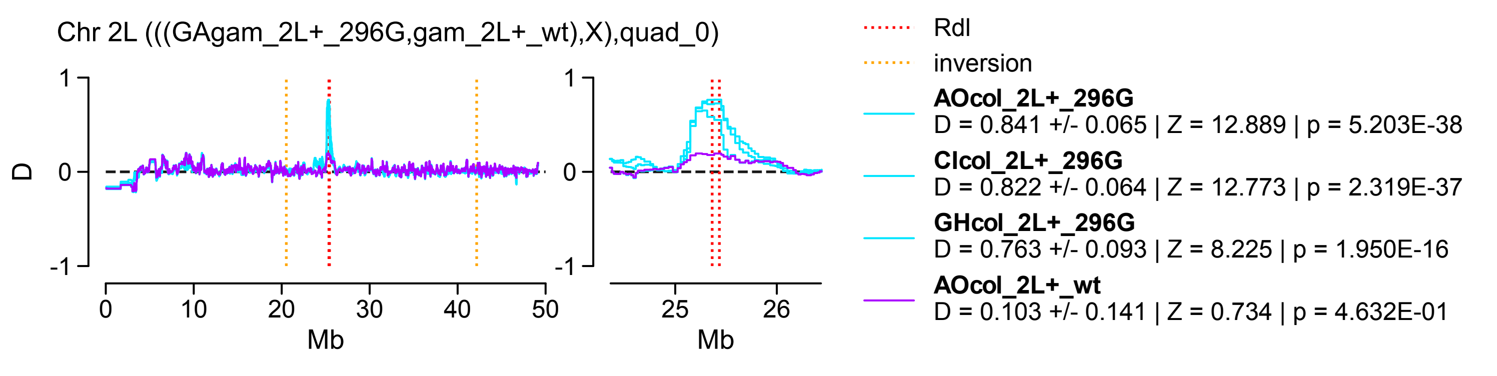


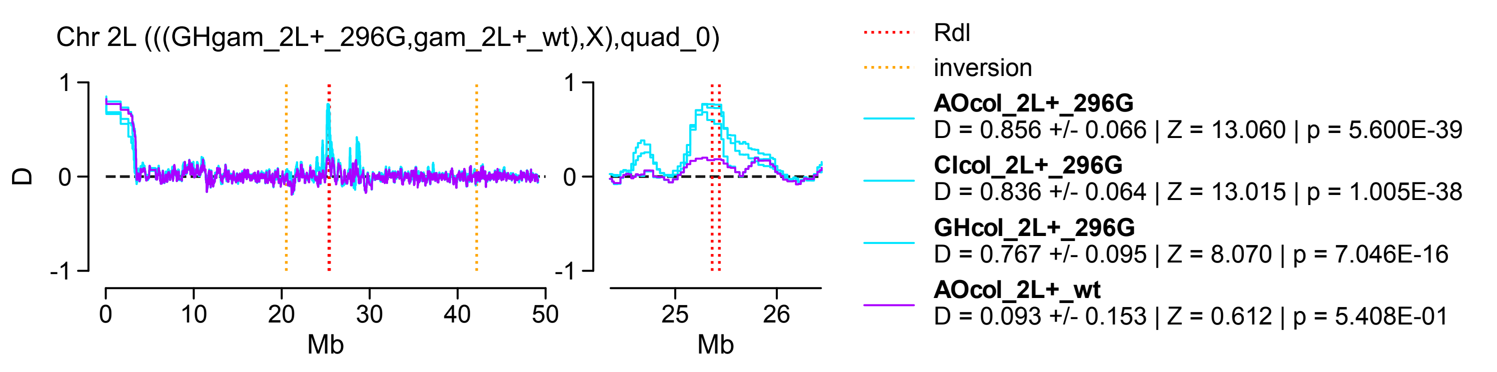


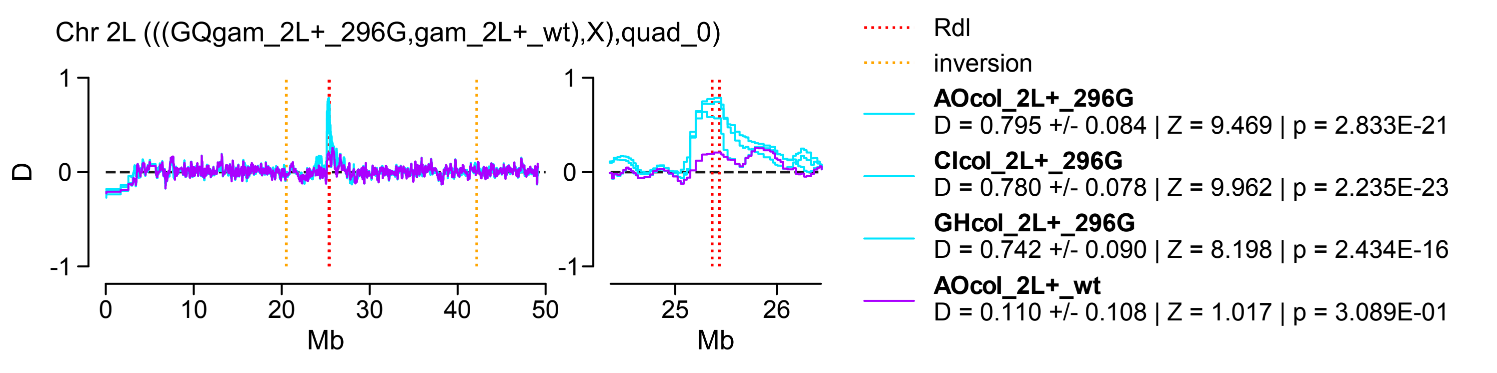


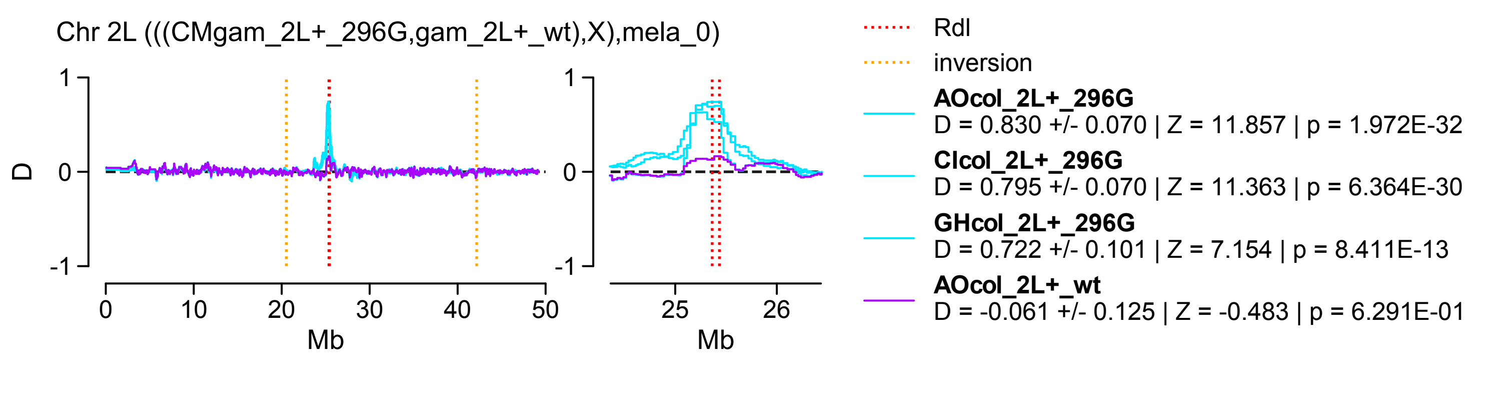


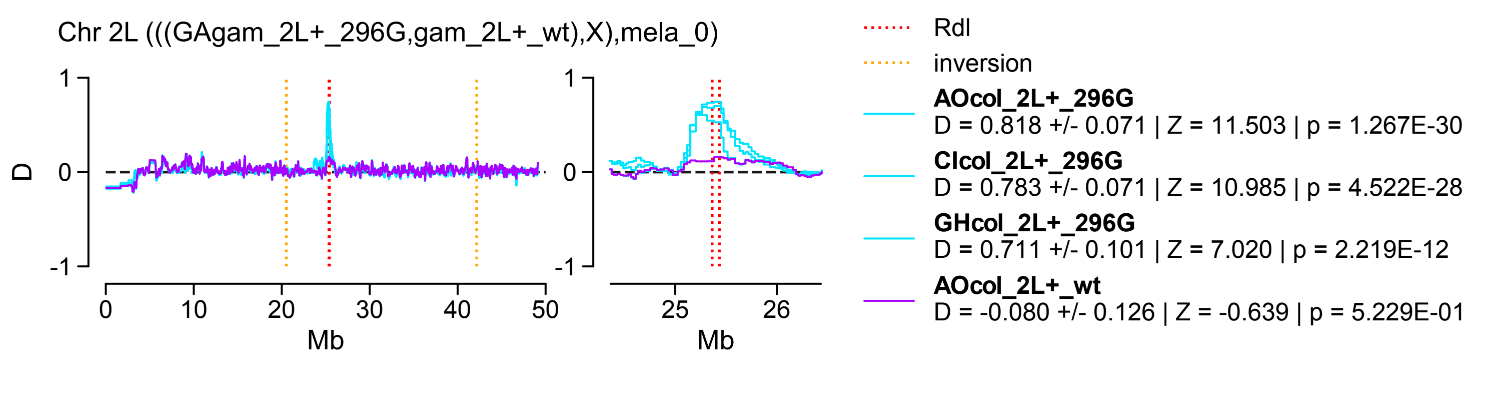


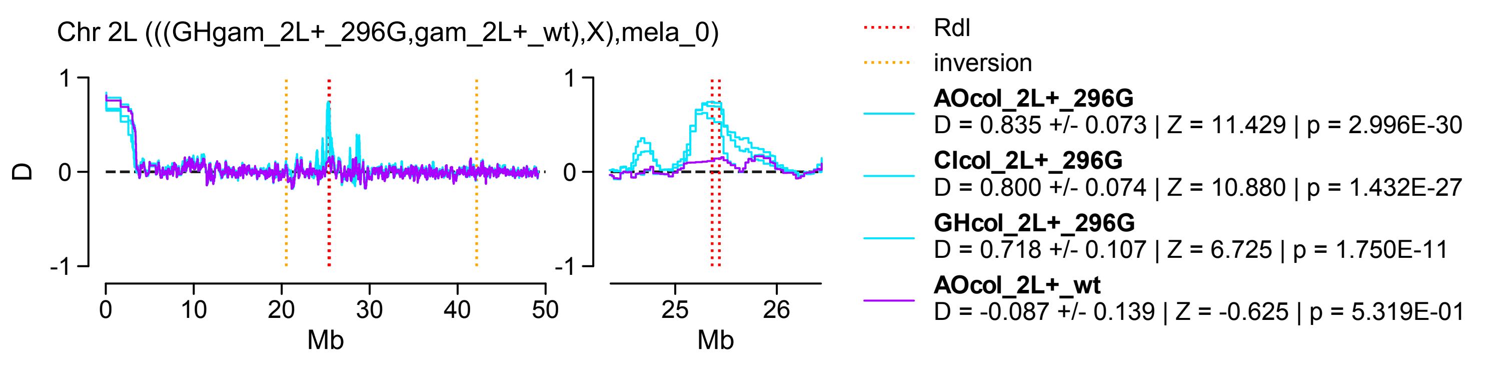


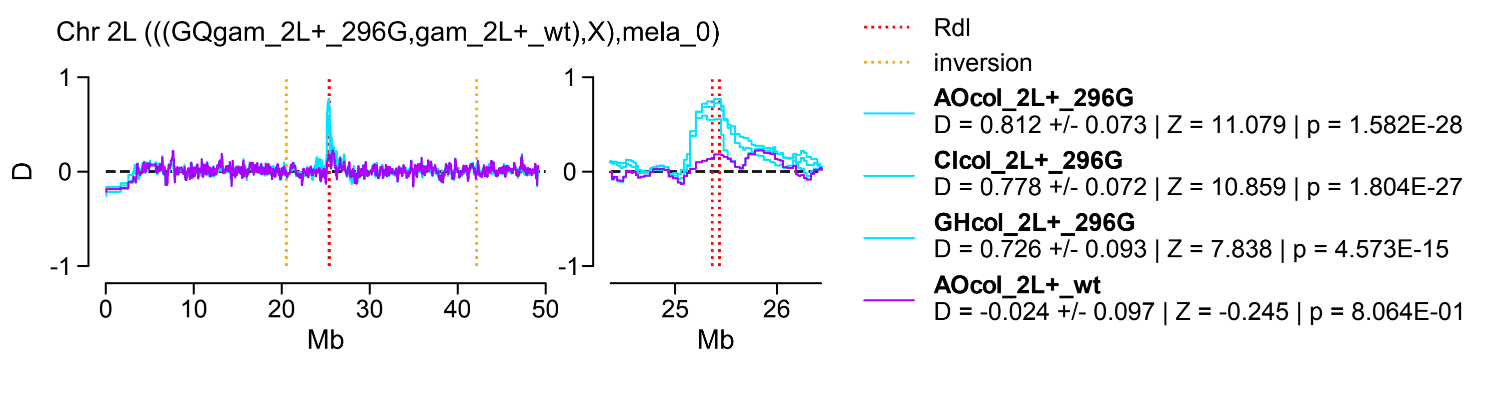


**C)** *A. gambiae* as donor (C), 2La homozygotes

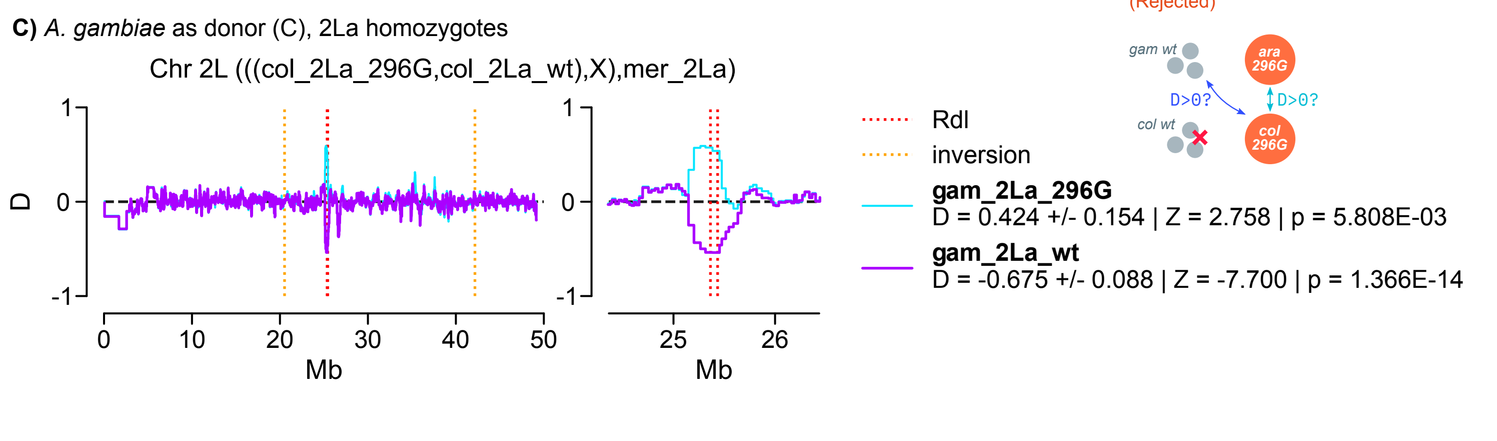


**D)** *A. coluzzii* as donor (C), 2La homozygotes

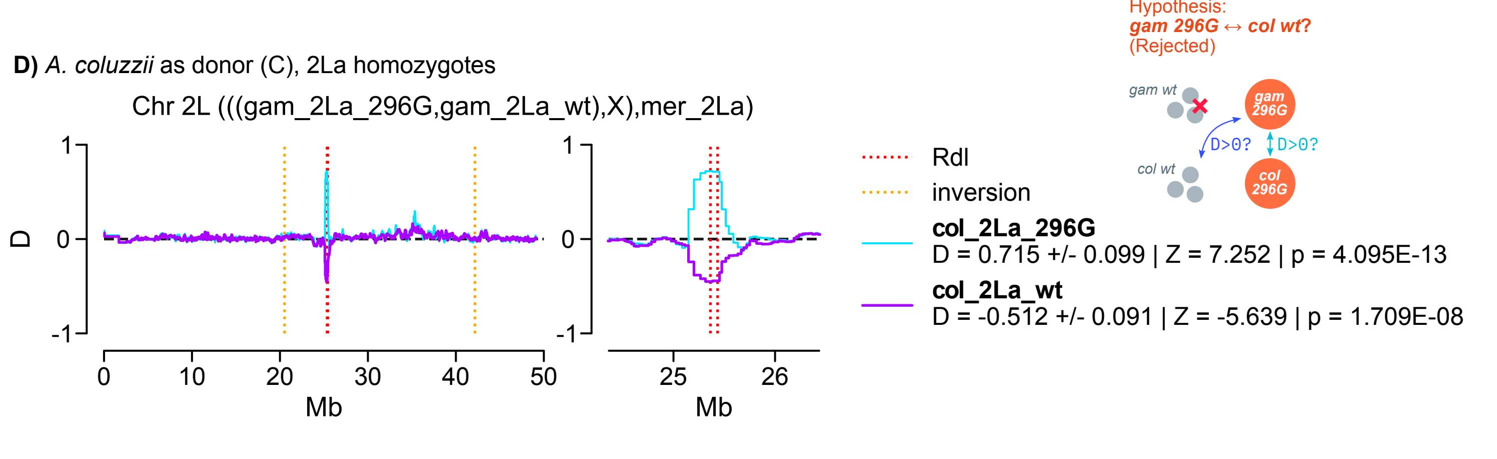
