## Supplementary Material SM14. Diversity of 296G haplotypes in 2L+a and 2La backgrounds for "Evolution of the insecticide target *Rdl* in African *Anopheles* is driven by interspecific and interkaryotypic introgression"

**Supplementary Material 14**

**A)** *EHH* decay 2L:25429236 +/- 150000, n=16623 vars

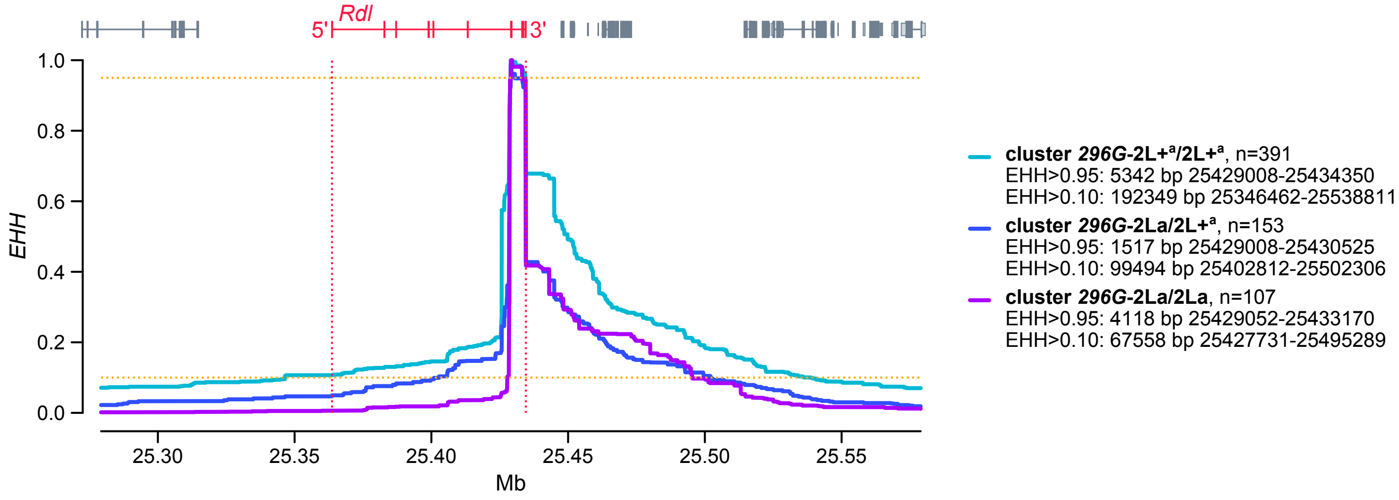


**B)** Haplotype diversity 2L:25429236 +/- 150000, n=16623 vars

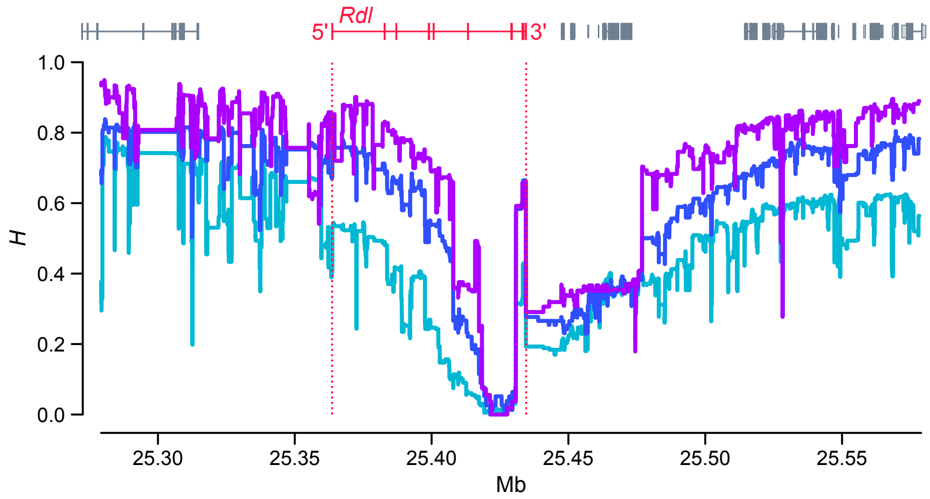


**C)** Sequence divergence between *296G* (2L+$^a$ background) and *wt* (2L+$^a$ or 2La) haplotypes

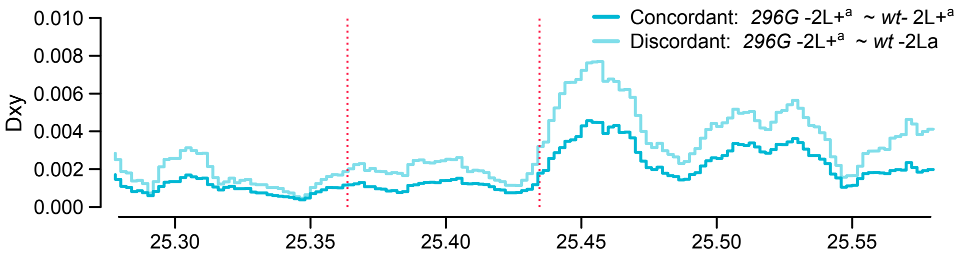


**D)** Sequence divergence between *296G* (2La background) and *wt* (2L+$^a$ or 2La) haplotypes

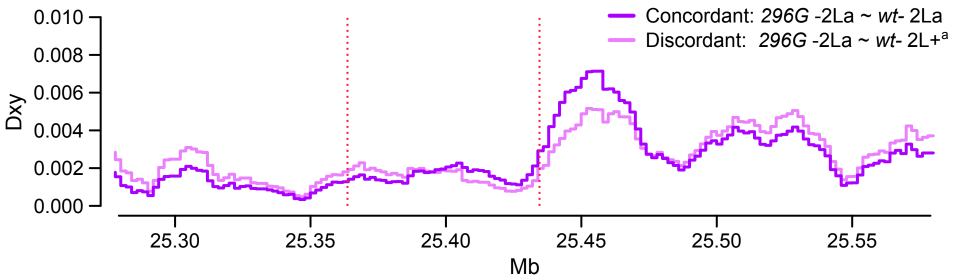
